# Distribution of genetic paternity in primate groups

**DOI:** 10.64898/2026.04.02.716091

**Authors:** Stacy Rosenbaum, Nicholas Grebe, Joan B. Silk

## Abstract

Understanding the distribution of paternity within social groups is critical for testing hypotheses about the evolution of behavior and morphology in primates, but assembling the requisite comparative data is a challenging task. We compiled genetic paternity data from 52 species of wild nonhuman primates along with information about socioecological, morphological, and life history traits that are relevant to understanding what proportion of offspring are sired by ‘primary males’ (i.e., alpha males in multi-male groups and resident males in single male groups). Our dataset, which currently contains information about 11 primate families and >3,000 individual paternities, is presented as a publicly accessible, living database designed to be updated as new data become available. Using Bayesian regression models, we investigated the role that phylogeny, group composition, and seasonality play in determining primary males’ paternity share, and assessed the relative share of paternities obtained by non-primary residents versus extra-group males. First, we found that phylogeny has a detectable but relatively modest influence on primary males’ paternity share. Species-level differences explained roughly 35–40% of variation in primary males’ paternity share, and of that interspecific variation, ∼50-70% was attributable to shared phylogenetic history. Second, group composition strongly predicted paternity share outcomes. Primary males in single-male/multi-female groups obtained the highest share of paternity (∼80%), while those in multi-male groups had the lowest (∼60%), though there was substantial variation within each category. Pair-living animals showed a striking split: males in cohesive pairs sired ∼90% of offspring, while those in dispersed pairs sired only ∼55%. Contrary to expectations, reproductive seasonality did not predict primary males’ paternity share in any group type. Finally, when primary males in multi-male groups lost paternities, ∼75% of losses were to other resident males. Overall, ∼5-15% of offspring in these groups were sired by extra-group males. Our results largely confirm earlier findings based on smaller datasets, but also show that the relationship between social organization and paternity is more complicated than simple categorical predictions suggest. We discuss the gap between the data that would ideally be available for testing these hypotheses versus what currently exists, with hopes that our living database can help close this gap over time.

## Introduction

Understanding the distribution of paternity within social groups or populations is critical for testing a wide variety of hypotheses about the evolutionary forces that shape male reproductive behavior and morphology. For example, incomplete control models of male reproductive skew predict that the extent of reproductive skew within groups will decrease with increases in the degree of reproductive synchrony among females and the number of males in groups (Ostner, Nunn, and Schülke 2008). The degree of sexual dimorphism that occurs in a given species is expected to be a product of the degree of male reproductive skew; in species where skew is high, sexual dimorphism in body size is greater and males are more likely to have ‘weaponry’ useful for fighting for access to females (Andersson 1994). In species that typically live in pairs, male care and the strength of pair bonds are expected to be inversely related to rates of extra-pair paternity (Huck et al. 2014). And, the evolution of cooperative breeding in mammals is expected to be linked to high rates of maternal and paternal relatedness within groups (Lukas and Clutton-Brock 2012).

Nonhuman primates are especially useful for exploring the drivers of paternity distribution because they are the second-largest mammalian Order and have extensive diversity in social organization and mating systems. The advent of DNA fingerprinting in the 1980s and the development of efficient methods for extracting DNA from fecal samples made it feasible to assess paternity in wild primate populations without capturing or handling them. The number of studies providing paternity data from wild nonhuman primates has steadily increased over time (Figure 1), and genetic paternity data are now available for more than 50 species. This represents a substantial increase over the number of nonhuman primate species that were available in previous reviews and comparative analyses (e.g., Clutton-Brock and Isvaran 2006; Widdig 2007; Ostner, Nunn, and Schülke 2008; Soulsbury 2010; Gogarten and Koenig 2013; Miller et al. 2021). In addition to increasing the total number of species available, researchers have accumulated larger samples of paternities for particular species and groups. Bigger sample sizes are particularly important if there is variation in patterns of paternity distribution across time within groups, or across populations within species; evidence from some nonhuman primate species suggests that this is frequently the case (Alberts, Watts, and Altmann 2003; Rosenbaum et al. 2015).

**Figure 1:**
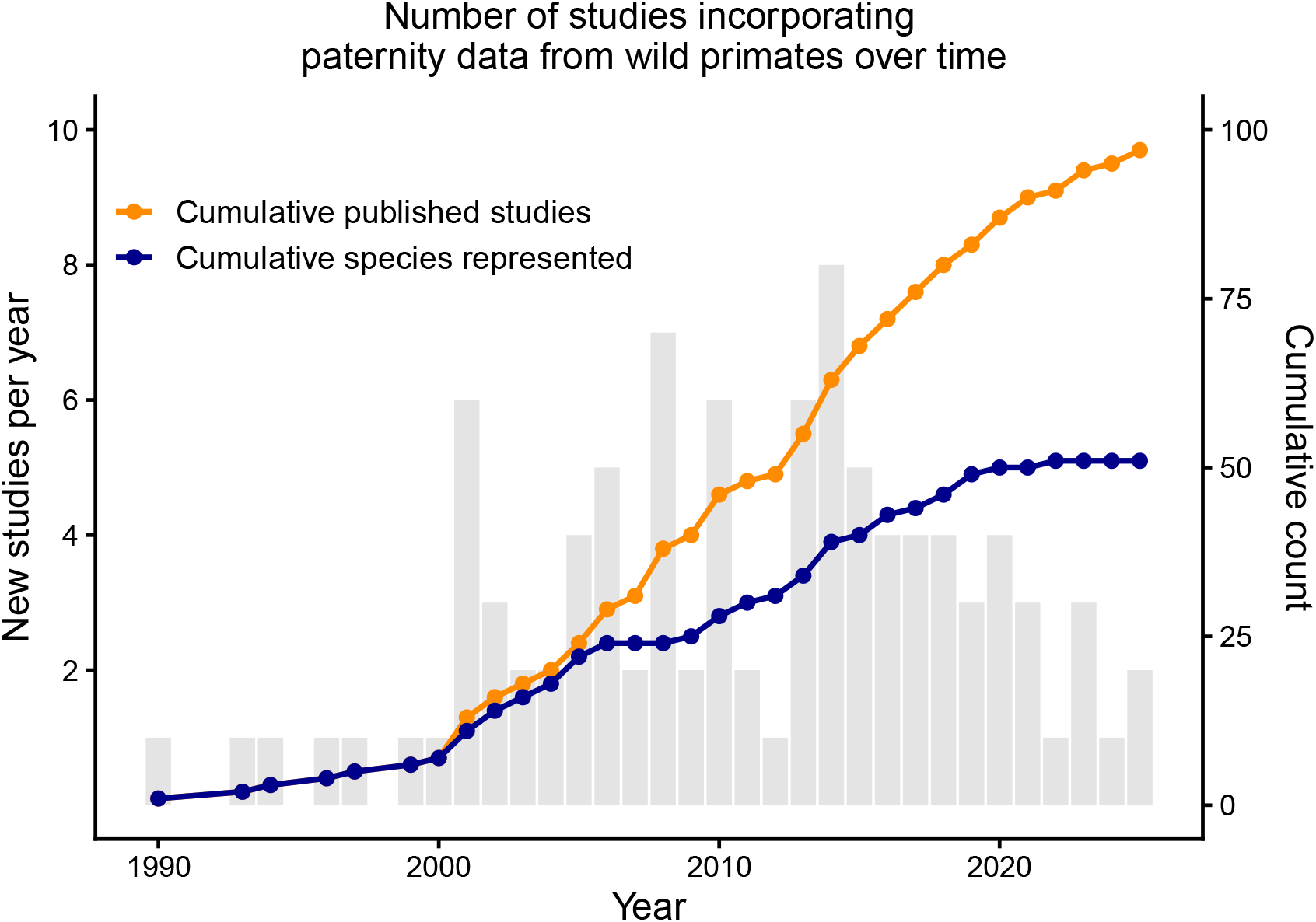
Grey bars represent the number of studies published each year between 1990 and 2025 that incorporate paternity data from wild non-human primates. The orange line tracks the cumulative number of studies over time, while the blue line tracks the cumulative number of species represented across those studies. Data in our table that are classified as personal communication/unpublished data are not included in this figure. Only one species, *Papio anubis*, was represented in unpublished data but not published data.

Compiling information about paternity distribution and phenotypic traits from primary sources is a labor-intensive task, making publicly available, compiled databases a valuable resource for the academic community. Comparative analyses are more powerful when they include information from more species, and they are more accurate when measurements of traits are based on larger and more representative samples.

We have compiled information about the distribution of paternity in wild primate groups, along with complementary information about socioecological, morphological, and behavioral characteristics that are potentially useful for testing evolutionary hypotheses about paternity distribution. A critical goal of our project is to compile all available information on paternity distribution in wild primate groups, make these data freely available to the academic community, and to create a mechanism for updating the database as new data become available. We attempt to achieve three key objectives for comparative datasets: transparency, usability, and reproducibility (Borries et al. 2016).

Another immediate goal of our project is to use these data to answer three basic questions about paternity distribution in non-human primates: first, to what degree does phylogeny predict paternities obtained by ‘primary males’—that is, alpha males in multi-male groups, and resident males in single-male groups? The answer to this question is useful for understanding the relative importance of evolutionary history versus current environmental conditions in determining paternity distribution.

Second, we evaluate the magnitude of paternity loss to other males for (a) resident males in pair-living groups, resident males in single-male/multi-female groups, and (c) top-ranked (alpha) males in multi-male groups. This is a test of the thus-far broadly supported prediction that paternity loss will be a function of group composition, where pair-living (i.e., socially monogamous) males have the highest paternity certainty and those in multi-male/multi-female groups the least (Clutton-Brock and Isvaran 2006). We also investigate whether the proportion of paternities obtained is influenced by reproductive seasonality. This is a test of the prediction that males in species where female fertility is highly synchronized will have more difficulty monopolizing mating opportunities, for which some prior meta-analyses have found evidence (e.g., Paul 1997; Ostner, Nunn, and Schülke 2008) and some have not (e.g., Kutsukake and Nunn 2006). Understanding the magnitude of group composition and seasonality effects is necessary to reconstruct the selective pressures shaping male reproductive strategies.

Third, we ask: when alpha males lose paternities in multi-male groups, do they lose them to residents or to extra-group males? This is useful for understanding the source of male-male competition, and the tradeoffs males make when they are either forced to, or choose to, live in social groups with other males.

## Methods

### Data compilation

#### Sources of information

Starting in 2020, JBS and SR began compiling a list of relevant studies from Google Scholar searches using key words that included: paternity, male reproductive success, reproductive skew, mating system, inbreeding, kin discrimination, and kin recognition. We also tracked citations from lists previously compiled by others to identify papers we did not find using our search strategy (e.g., Widdig 2007; Ostner, Nunn, and Schülke 2008; Baker and Shackelford 2018; Miller et al. 2021). We almost always obtained quantities from the original publications, not from secondary sources; exceptions are noted with full references provided in our database. For published abstracts that contained insufficient information on their own, we contacted lead authors and requested additional details. In these cases, the abstract is cited along with a notation that more detailed data were obtained via personal communication. We also contacted researchers that we judged likely to have unpublished paternity data and requested any information that they were willing to share.

Some studies provide data from multiple social groups in the same population. When possible, we present information for each group separately. In some cases, these group-level data were available in the original publication. In other cases, we obtained information about individual groups from the authors of the original publication. In these cases, we cite the original source, along with a notation that additional information was obtained through personal communication with the authors.

In some instances there were multiple published studies from the same group of primates and the paternity data overlaped (e.g., the capuchin paternities in Muniz et al. (2010) are a subset of the paternities in Godoy, Vigilant, and Perry (2016)). In these cases, all the studies are included in our dataset, and their overlap is indicated along with information about which study contains the largest sample size. To avoid duplication of data, our analyses are based on the study with the largest sample size. We provide a full bibliography listing all included studies in the supplementary materials.

#### Types of information

Below we describe the data we aggregated, loosely subdivided into types of information for organizational purposes. We combined information from paternity studies with information from other studies of the relevant species. The data are arranged so that each row corresponds to a particular social group (whenever possible) or to an aggregation of multiple social groups (when it was not possible to extract group-level statistics from a given publication).

#### Study identification

To identify individual studies and track entries over time, we include the reference to the paper the data came from, the year the paper was published, the data source (i.e., whether it was primary data or secondary), and the date we added the study to our table. These variables are listed in Table 1.

**Table 1.** Study identification variables.

| Variable name | Variable description |
| --- | --- |
| reference | Study the data came from |
| publicationyear | Publication year |
| datasource | Primary or secondary data source; if secondary, refs in notes |
| dateofentry | Date that study was entered into the database |

#### Study context and genetic sample characteristics

Variables that contain basic information about the study context and genetic data are listed in Table 2. We provide information about the location and name of the field site, whether the animals were provisioned or not (in n=10 studies on 4 species the animals were provisioned or commensal in some way), the years in which paternity data were collected, what sample matrix or matrices the DNA was obtained from, the number of loci that were evaluated, the paternity confidence levels that were used, and whether the analyses were primarily longitudinal or cross-sectional. As described above, we also noted whether there was known data overlap with other studies, and if so, which of the studies was the largest/newest. In all cases, the newest publication also contained the largest number of paternities.

**Table 2.** Study context and genetic sample characteristics.

| Variable name | Variable description |
| --- | --- |
| country | Country where the study was conducted |
| field_site | Name of the field site |
| condition | Whether the population was provisioned or wild-feeding |
| year_first | Earliest year included in study |
| year_last | Latest year included in study |
| n_yrs_sampled | Number of years sampled |
| sample_matrix_1<sup>1</sup> | Type of matrix used (e.g., feces, tissue) |
| loci_lo | Minimum number of loci used |
| loci_hi | Maximum number of loci used |
| paternity_conf_strict | Paternity confidence level (strict) |
| paternity_conf_relax | Paternity confidence level (relaxed) |
| long_or_cross | Whether analysis was primarily cross-sectional or longitudinal |
| study_overlap | Do paternity data in this study overlap with another study? |
| study_overlap_name | If there is overlap with another study, which one? |
| newest_biggest | If there is overlap with another study, is this study the newest/biggest? |

<sup>1</sup> Actual table contains sample\_matrix\_1, sample\_matrix\_2, etc. up to sample\_matrix\_6, because some studies include DNA from multiple sources (e.g., feces and tissue).

#### Study subject characteristics

Study subject variables are listed in Table 3. We included the common and scientific species name provided by the authors of the study, as well as the genus, species, and (where applicable) subspecies name used by the 10kTrees website (Arnold, Matthews, and Nunn 2010). We used this website to generate the distance matrix needed for our phylogenetically informed analyses. In the vast majority of cases these names were identical.^1^

**Table 3.**
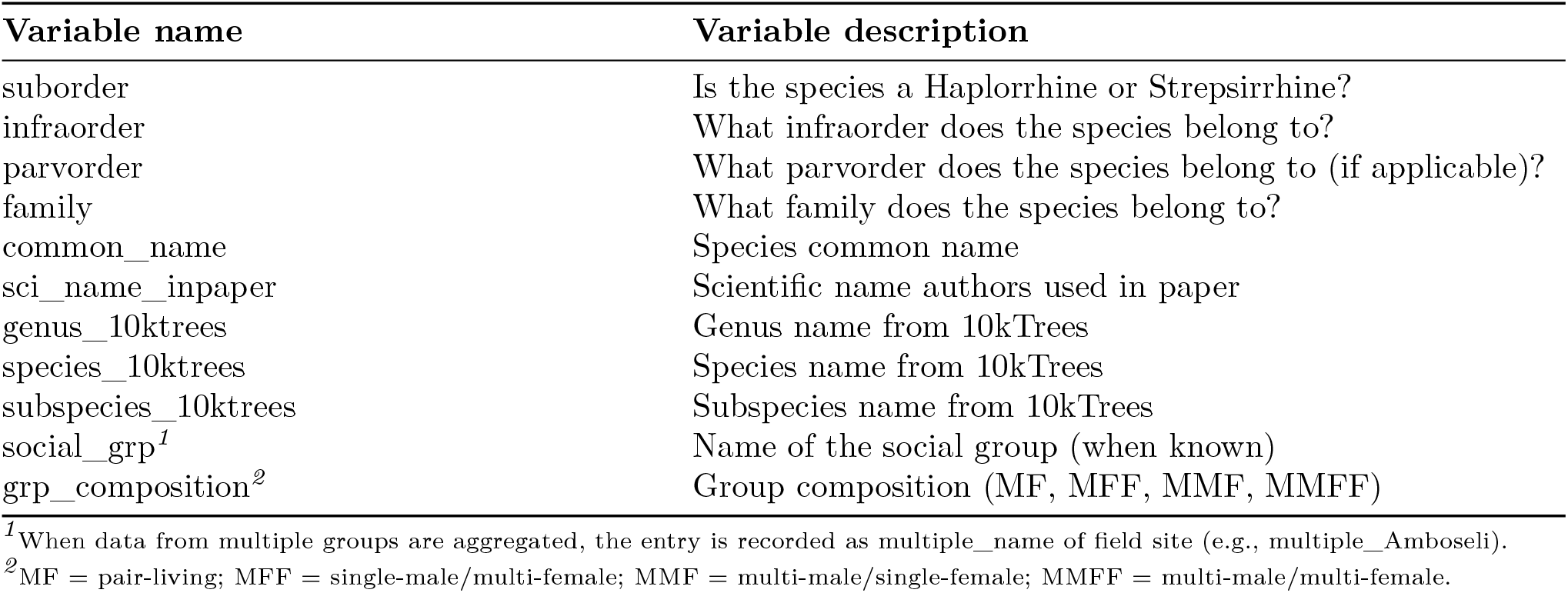
Study subject characteristics.

| Variable name | Variable description |
| --- | --- |
| suborder | Is the species a Haplorrhine or Strepsirrhine? |
| infraorder | What infraorder does the species belong to? |
| parvorder | What parvorder does the species belong to (if applicable)? |
| family | What family does the species belong to? |
| common_name | Species common name |
| sci_name_inpaper | Scientific name authors used in paper |
| genus_10ktrees | Genus name from 10kTrees |
| species_10ktrees | Species name from 10kTrees |
| subspecies_10ktrees | Subspecies name from 10kTrees |
| social_grp<sup>1</sup> | Name of the social group (when known) |
| grp_composition<sup>2</sup> | Group composition (MF, MFF, MMF, MMFF) |

<sup>1</sup>When data from multiple groups are aggregated, the entry is recorded as multiple\_name of field site (e.g., multiple\_Amboseli).
<sup>2</sup>MF = pair-living; MFF = single-male/multi-female; MMF = multi-male/single-female; MMFF = multi-male/multi-female.

We also recorded the names of the social groups whenever they were provided. In cases where numbers from multiple groups were aggregated, we recorded the group name as multiple_ followed by the name of the field site (e.g., multiple_Amboseli). We also classified groups based on the number of adult residents of each sex living together as a cohesive unit, rather than the presumed mating system for the taxa. Group composition was categorized as follows: pair-living = one resident (adult) male and one resident (adult) female; single-male/multi-female = one resident male and two or more resident females; multi-male/single-female = two or more resident males and one resident female; and multi-male/multi-female = two or more resident adults of each sex. The abbreviations MF, MFF, MMF, and MMFF, respectively, are used in the actual dataset as shorthand (Table 3). In some cases, group composition changed across time, so we tabulated data separately for each group composition type whenever possible.

#### Information about paternity distribution

Paternity distribution measures are listed in Table 4. Authors use different metrics to report paternity results, but for the majority of studies it is possible to tabulate the total number of paternities tested along with (a) the number of paternities obtained by the resident male in groups that include only one male, or (b) the number of paternities obtained by the top-ranking (alpha) male in groups that include more than one male. We categorized (a) and (b) together as primary males. We also extracted the number sired by other group residents of 1) known rank and 2) unknown rank. Note that the second category includes studies where the rank(s) of the males in question were unknown due to lack of behavioral data, where they were not reported, or where the authors were unable to conclusively assign a single in-group male as the father. In addition, we extracted the number of individuals that were sampled but for whom paternity remained undetermined, number of extra-group paternities, number where whether the paternity was extra-group was ambiguous, number sired by the single most successful male in the study, and number sired by all other (i.e., not most successful) males combined. These categories are not necessarily mutually exclusive, so the summed totals across rows may not match the total number of individuals that were sampled. For example, a given paternity could appear in both the column that tabulates the number of individuals for whom paternity was undetermined and in the column that tabulates instances where extra-group paternity was ambiguous, because the authors identified two equally likely potential sires but did not have demographic data to determine whether these potential sires were residents of the group at the time of conception.

**Table 4.** Paternity distribution information.

| Variable name | Variable description |
| --- | --- |
| n_paternities | Number of paternities tested |
| n_alphaprimary | Number of paternities attributed to the alpha or resident male |
| n_residents_knownrank | Number of paternities attributed to resident males of known rank |
| n_residents_unknownrank | Number of paternities attributed to resident males of unknown rank |
| n_tested_undetermined | Number of paternities that were tested but specific sire was not determined |
| n_egp | Number of established extra-group paternities (EGPs) |
| n_egp_ambiguous | Number of paternities that cannot be confidently assigned as either EGPs or resident sirings |
| n_sired_mostsuccess | Number of paternities attributed to the most successful male |
| n_sired_othermales | Number of paternities attributed to all other (not most successful) males |
| mean_sire_rank | Mean dominance rank of sires, when ranks are known |
| meanrank_standard | Is mean rank of sires standardized (Y/N)? |
| nonacs_b<sup>I</sup> | Nonacs' B index of reproductive skew |
| ciforbindex_low | Lower confidence interval for Nonacs' B |
| ciforbindex_high | Upper confidence interval for Nonacs' B |
| nonacsb_p | p-value for Nonacs' B |

<sup>I</sup>When a study contained Nonacs' B-index values from >1 point in time, these are recorded as nonacs\_b\_t2 and nonacs\_b\_t3. The same applies to the ciforbindex\_low and \_high entries (which contain the low and high confidence intervals associated with a B-index at t, t2, t3), and to the nonacsb\_p entries (which contain the p values associated with the B-indexes).

In addition to count data, some studies report relevant metrics such as the mean rank of sires or the B-index of reproductive skew (Nonacs 2000). While Nonacs’ B-index (NB) suffers from flaws (namely, increasing values in larger group sizes are a statistical artifact of the way the index is constructed (Kutsukake and Nunn 2006; Ross et al. 2020)), it remains the most commonly reported skew index value in the primate literature. We did not recalculate NB or mean rank of sires if they were not provided by the authors, because the necessary data to make these calculations were often unavailable. In cases where multiple NB values were included in a single study (e.g., at multiple points in time), we recorded all available NB values, confidence intervals, and p-values. When known, we also recorded whether or not the authors had genetic data for all of the resident males.

No single study had all of these metrics available, and some had only a small number of them. When we had meaningful doubt about our ability to accurately calculate or extract a desired metric, we emailed the first author of the study to request clarification. In some cases they did not respond, in which case we left out a number that in theory could have been calculated. If the authors of any of our included studies find mistakes or fillable gaps in the table, we welcome the chance to update our existing information and to add new information. For studies with so little extractable information that they were not included in any of the present analyses, we still recorded them in the interest of maintaining a complete list of studies that have primate genetic paternity data.

#### Information about the species

Species-level socioecological, morphological, and life history information is listed in Table 5. This was almost always obtained from sources other than the genetic paternity studies in our table; much of it came from Kamilar and Cooper (2013) and Lüpold, Simmons, and Grueter (2019). Not all measures were available for all species. Many of the available measures rely on older data or small sample sizes. If experts are able to provide more up-to-date species-level information, we welcome the opportunity to improve our dataset.

**Table 5.** Socioecological, morphological, and life history traits.

| Variable name | Variable description |
| --- | --- |
| seasonality_5c<sup>*I</sup> | Degree of birth seasonality in natural habitat (1=most, 5=none) |
| seasonality_5c_2<sup>*I</sup> | Degree of birth seasonality in natural habitat (1=most, 5=none) |
| seasonality_3c<sup>*I</sup> | Degree of birth seasonality in natural habitat (1=most, 3=none) |
| multilevel | Does species live in multilevel society? |
| philopatry_pattern | Primarily male or female philopatric? |
| pairbond_type | In pair-living groups, are pairs dispersed or cohesive? |
| gestation_length | Gestation length (days) |
| testes_mass | Combined weight of testicles (g) |
| litter_size | Mean litter size |
| mean_rainfall_mm | Mean precipitation (mm) |
| mean_male_mass | Average male weight (g) |
| mean_female_mass | Average female weight (g) |
| mass_dimorphism | Weight dimorphism value |
| canine_dimorphism | Canine dimorphism value (canine length) |
| mean_grp_size | Average size of social group |
| mean_num_males | Average number of males who live in a social group |
| mean_num_females | Average number of females who live in a social group |
| adult_sex_ratio | Ratio of adult females to adult males |

<sup>I</sup>Birth seasonality classifications were obtained from Heldstab (2021). Heldstab includes three different classification schemes; our dataset includes all three.

#### Frequency of extra-group paternities (EGPs)

Many studies directly report the number of paternities obtained by males from outside the group (extra-group paternities, EGP). In some cases, paternity was assigned to specific extra-group males who had been sampled. In other cases, all resident males were sampled and could be excluded from paternity, and we thus assigned these paternities to extra-group males. If not all potential sires from within the group were sampled, paternities that could not be assigned to known male residents were categorized as ambiguous (variable n_egp_ambiguous).

#### Proportion of conceptions by primary males and extra-group males

For cases in which all paternities could be assigned to primary males, other resident males, or extra-group males, the exact proportions of paternities by the primary male, other residents, and extra-group males were calculated by dividing the number of paternities in each category by the sum of paternities by primary males, other residents, and extra-group males.

When paternities were categorized as residents of unknown rank, we calculated minimum and maximum estimates obtained by primary males. For the minimum estimate, we assumed that none of the paternities attributable to residents of unknown rank were sired by primary males. We divided this number of paternities by the sum of all paternities (i.e., infants sired by the primary male, other residents, EGPs, and cases where it was ambiguous whether it was an EGP or in-group siring). To calculate the estimate of the maximum proportion sired by primary males, we assumed that all paternities in this category were sired by primary males, and divided by the same sum.

Similarly, in cases in which some paternities were categorized as ambiguous, we calculated minimum and maximum estimates of the proportion of paternities by extra-group males. To calculate the minimum estimate, we assumed that all paternities in this category were sired by resident males, and divided the number of paternities by extra-group males by the sum of total paternities. To calculate the maximum estimate, we assumed that all paternities in this category were sired by extra-group males, and divided by the same sum. The true proportion lies somewhere in between these two values.

In practice, to whom uncertain paternities were assigned made little difference to our qualitative conclusions. Unless otherwise noted, when proportion of total paternities is the target outcome, we use primary males’ maximum possible paternity share. When count of EGPs is the outcome, we use only verified EGPs (the most conservative estimate). The supplementary materials contain results from models that use minimum possible primary male share and maximum possible EGPs for comparison purposes (Tables S2-S13).

#### Theoretical model

To clarify what data would ideally be needed to test the relationship between competition and paternity, and to make explicit the assumptions that underlie our empirical approach, we developed a simple formal theoretical framework. We assume that the proportion of paternities (*p*) that a primary male in a social group will obtain is, at its most basic, a function of two things: first, the number of competitors that he has (*n*), and second, the degree to which the mating opportunities in his group are monopolizable (*m*) (e.g., Kutsukake and Nunn 2006). In other words:

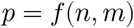

We assume that as *n* increases, *p* should decrease (because more competitors likely equals more paternities lost), and that as *m* increases, *p* should increase (because greater monopolizability should mean more paternities gained). A simple model of this relationship could be:

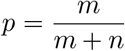

Figure 2 contains a visualization of these dynamics. This is in most respects simply Altmann’s priority-of-access model (Altmann 1962), but it 1) focuses specifically on the paternity share of one animal (i.e., the primary male), 2) replaces the number of estrus females with fertile female:male ratio, and 3) makes no assumptions about the competitive abilities of individual males.

**Figure 2:**
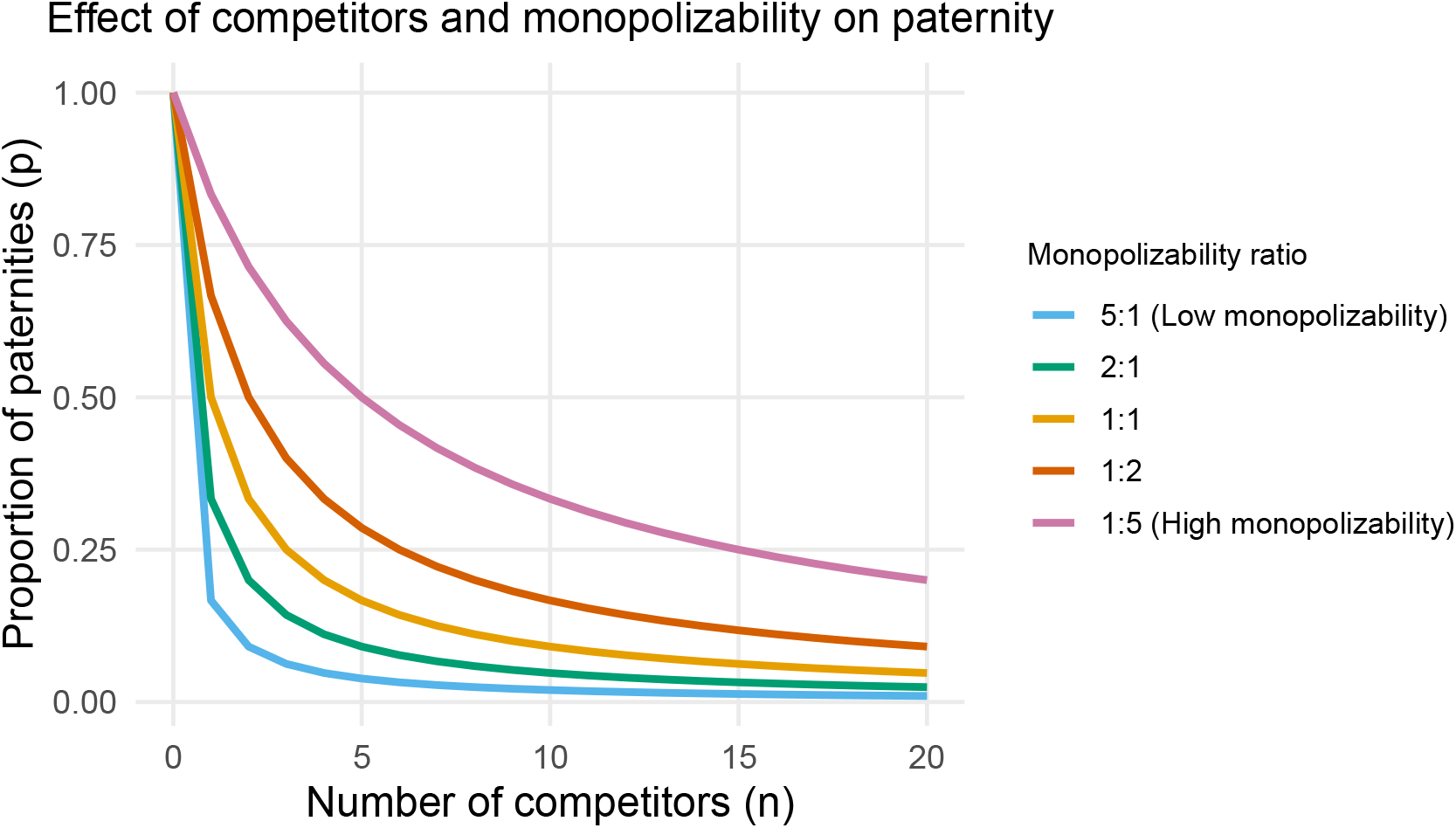
Expected change in the proportion of paternities obtained by primary males as a function of number of competitors and fertile female:male ratio. E.g., a 5:1 ratio (the blue line) would represent a situation where a large number of females were fertile at the same time, and thus monopolizability is low (for example, in a seasonally breeding monkey species). A 1:5 ratio (represented by the purple line) would represent a situation where individual females only occasionally come into estrus (for example, a chimpanzee group that contains many males but likely only one or at most two fertile females at any given point in time), and so monopolizability is high.

Since competition can come from either males within the same social group or males from outside of it, *n* can be further distinguished as follows:

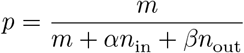

where *n*_in_ is the number of within-social group competitors a male faces, *n*_out_ is the number of extra-group competitors he faces, and *α* and *β* are weighting parameters that indicate the relative strength of intra- and extra-group competitors (e.g., in mountain gorillas, extra-group competitors may be harder to drive off, perhaps because they are more motivated to fight for mating opportunities (Robbins and Sawyer 2007)).

The value of this theoretical framework is that it makes clear precisely what data would be needed for a rigorous test. Specifically, we would need: (1) conception-level data on social conditions at the time each offspring was sired, rather than paternity proportions aggregated over long time periods; (2) a continuous measure of the number of competitors, including some way to separately quantify within-group (*n*_in_) and extra-group (*n*_out_) competition; and (3) a direct measure of monopolizability at the time of conception, rather than a species-level proxy.

Few papers in our dataset report this information. Due to primates’ slow life histories and the realities of sample collection and analysis, paternity determination usually does not happen until months or even years after offspring are born. The social conditions of the father at the time of conception of a given offspring are frequently omitted from studies, sometimes because they are not available, and sometimes because they are not seen as relevant. Small sample sizes mean that data are often aggregated over many years. However, unless group composition and number of mating opportunities are extremely stable, statistics like the proportion of infants sired by primary male(s) across long time periods hide the potentially considerable amount of variation in social conditions that may have occurred at the point of the conception, which is the biologically relevant time window.

Given these constraints, our empirical analyses depart from the theoretical model in several ways. We operationalized the number of competitors as a binary indicator of group composition (single-male vs. multi-male), which collapses the (ideally) continuous *n*_in_ term to a categorical variable and leaves *n*_out_ unmeasured. For monopolizability, we used a species-level index of reproductive seasonality (Heldstab et al. 2021). This assumes that *m* is a reasonable proxy for the temporal distribution of fertile periods, a potentially questionable assumption (see, e.g., Ostner, Nunn, and Schülke 2008; Gogarten and Koenig 2013) but also ignores other potentially important factors such as female spatial distribution.

While these metrics are much less precise than individual conception-level information would be, they are reasonable proxies given the constraints of currently available data. The theoretical model helps make these compromises and their implications transparent. More details on our empirical implementation are provided below in the data analysis section.

#### Data analysis

All analyses were run in R 4.3.2 and all models were fit using brms (Bürkner 2017), a flexible software package for fitting Bayesian multilevel models with a variety of response distributions, link functions, and prespecified correlation structures. These features make brms well-suited to our phylogenetic analyses of paternity patterns. We constructed a phylogenetic covariance matrix using the consensus primate tree available from 10kTrees (Arnold, Matthews, and Nunn 2010) that is included in all models below as a random effect with a covariance structure corresponding to evolutionary distances between species (see Bürkner (2025) for a vignette of these models in brms).

The proportion of paternity for a given class of males (i.e., primary, resident, extra-group) can span the entire range from 0 to 1. While Beta regression is a popular option to model proportions as response variables, it cannot accommodate proportions at either extreme–i.e., exactly 0 or exactly 1, a situation which is common in our dataset. While there are modifications that can accommodate values of exactly 0 and/or 1 (e.g., zero-one inflated Beta: Liu and Kong (2015)), an underlying assumption of such models is that the processes that produce zeros, ones, or both are different in some way than the processes that produce the rest of the distribution. We do not think that this is a biologically defensible stance for our analysis. The fundamental processes that produce zeros and ones are likely the same as the ones that produce less extreme values. Furthermore, *true* zeros and ones are probably unlikely to occur; it seems implausible that there are species in the real world where primary males either obtain every possible paternity or never obtain any of them. It is more likely that our 0/1 values are the result of small sample sizes and the potential for biased sampling of certain paternities. Thus, in order to use a standard Beta regression model, we replaced observed proportions of 1 with 0.999, and observed proportions of 0 with 0.001.

For our first analysis examining the influence of phylogeny alone (Question 1 in the results), we specified our models predicting our various operationalizations of paternity proportions as:

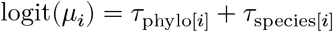

where the observation-level paternity proportions (*μ*_*i*_) are predicted by random effects (*τ*) for phylogeny and species. The *τ*_phylo[*i*]_ term captures variation due to phylogenetic relatedness between species, and its variance quantifies the amount of interspecies variation that is attributable to phylogeny. The *τ*_species_term estimates interspecific variance not explained by phylogenetic relatedness. From these estimates, we calculated the phylogenetic signal (*λ*) by dividing the total variance from phylogeny 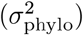 by the sum of the variance sattributable to phylogeny plus the variance attributable to species-specific variation that is not explained by phylogeny:

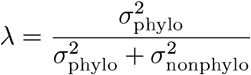

Thus, *λ* is the proportion of interspecies variance that is phylogenetically structured. Another way to state this is that it answers the question “of the variation in primary male paternity share between species, how much of it is explained by phylogeny?” This is analogous but not identical to Pagel’s *λ* (Pagel 1999), which is estimated by directly transforming the phylogenetic tree to find the best fit to the data and incorporates total residual variance. Unlike Pagel’s *λ*, our *λ* is a ratio of the variance components derived from the model’s random effects, and excludes observation-level variance from the Beta distribution. We chose this approach because it directly addresses the between-species question and benefits from the observation weights and Beta likelihood included in the model.

Our second question is about the role that group composition (as a proxy for degree of male-male competition) and seasonality (as a proxy for monopolizability) play in primary males’ paternity share. To evaluate the effects of group composition and seasonality, first, we conducted a basic analysis that compared primary male paternity share values in multi-male, single-male, and pair-living groups; we present the raw data but also model-derived estimates that account for phylogeny and sample size (Analysis 1 in the results). Second, to determine how the combined influences of competition and seasonality affect paternity share (Analysis 2 in the results), we turned to our theoretical model described above (*p* = *m*/(*m* + *n*)). This model assumes a fixed functional relationship among the variables—specifically, that paternity share is determined by the ratio of monopolizability to competitive pressure exerted by other males. This is a useful conceptual foundation but may be too restrictive to capture the range of empirical variation observed across species. To allow for more flexibility, including potential non-linear effects of competitors and monopolizability, we constructed a model that contained main effects for both *n* and *m* plus an interaction between the two, as well as a squared term for *m*, while retaining the same random effects structure described above:

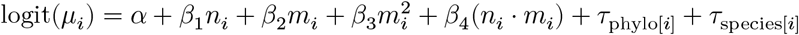

Because we were only able to operationalize competitors as a categorical variable where 0=no in-group competitors (i.e., males living in single-male groups) and 1=one or more in-group competitors (i.e., males living in multi-male groups), a squared term for *n* would be entirely redundant with the main effect term and is thus not included. Monopolizability was operationalized as a reproductive seasonality variable obtained from Heldstab et al. (2021). For this, lower values indicate more seasonality (and thus lower monopolizability) and higher values equal less seasonality (and thus higher monopolizability). To interpret the results we plotted the marginal effects and computed the partial derivative (∂) of *log*_*i*_*t*(*μ*_*i*_) with respect to monopolizability 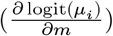, evaluated at the mean of *m* (i.e., moderate seasonality). For reporting purposes we transformed this quantity to the response scale via the chain rule, which provides the estimated change in paternity share per unit increase in monopolizability. For the binary competitor variable *n*, we interpreted the contrast in predicted values between single-male and multi-male groups.

Our third question was about the source(s) of primary male paternity loss in multi-male groups. To answer it, we fit a binomial generalized linear mixed model (GLMM). Here, the outcome variable is the number of EGPs for a given observation, treated as the number of “successes” out of total count of paternities not attributed to the primary male (i.e., EGPs plus sirings by other, non-primary resident males). Formally, we specified this as:

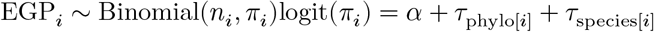

where EGP_*i*_ is the observed number of EGPs for observation _*i*_, *n*_*i*_ is the total number of non-alpha paternities for that observation, and *π*_*i*_ is the probability that a non-alpha paternity is attributable to an extra-group male. For this model we kept our zero-loss observations as zeros rather than rounding to 0.001, because in this case we were modeling counts (i.e., “0 trials out of X trials”) rather than a continuous distribution of probabilities. Consistent with the logic for our prior model specifications, zeros likely do not represent a fundamentally different biological sampling process than non-zeros, so we elected not to use (e.g.) a hurdle or zero-inflated model.

All Beta regression models were weighted by the square root of the total number of paternities in a given study, to assign greater weight to larger studies that had more precise estimates of paternity distribution proportions. For our binomial model (Question 3), no additional weighting was applied because the binomial likelihood already accounts for sample size through the number of trials. We set a Gaussian (normal) prior on the intercept, centered at 0 (SD=1) on the logit scale. This means we set a weak prior centered on primary males obtaining 50% of paternities, while allowing for significant uncertainty and a wide range of plausible values. For the standard deviation of the random effects and the distribution precision parameter (*ϕ*), we used Student’s t-distribution priors (df=3, center=0, scale=2.5). These are weakly informative, to allow for the possibility of substantial species-level variation and data dispersion, both of which we observed in our initial data exploration. We used the same strategy for the binomial model, except that the precision parameter was excluded. All models easily passed convergence diagnostics: Rhat values were close to 1.00 (well below the 1.01 threshold); bulk and tail effective sample sizes were substantially above recommended minimums; and there were no divergent transitions (Gelman et al. 1995; Vehtari et al. 2021). See Table S1 in the supplementary materials for details.

#### Living table and results

Across scientific disciplines, researchers have recognized the value of building “living” systematic reviews and/or databases. While there are numerous tools, techniques, and strategies for providing these resources, all of these approaches share a common goal of creating scientific products that facilitate mechanisms for update (see e.g., Elliott et al. 2017; Ben Mocha et al. 2024). In deciding how to structure the database and the results derived from it in a continually updateable format, we drew inspiration from the Cochrane Guidance for Living Systematic Reviews (Brooker, Synnot, and McDonald 2019). Below are the core features of our living resources:

1. The results reported below in this static manuscript are based on the version of the dataset as of March 2026.
2. Twice a year, we will perform a new search for relevant data, using the strategy described above in the data compilation section.
3. If new data are identified, the relevant features will be extracted, screened, and aggregated according to the methods described above.
4. We will provide the most current version of the dataset on an open-access Github repository (https://github.com/slrosen/primate-paternity-metanalysis). All updates to the underlying dataset will be documented in a README file.
5. The most current version of our results, plus all previous versions, will be posted on the same Github repository.

## Results

### Dataset description

Our database contains 264 entries from 52 species representing 31 genera and 11 families, encompassing a total of 3,063 unique paternities. Of these entries, 59 were excluded from the current analyses either because of data redundancies or because they were irrelevant for our questions, while an additional 57 lacked sufficient information to calculate primary male paternity share (see Table 6). This left 148 entries, 46 species, and 2,432 paternities for the analyses presented below. The phylogenetic distribution of the species in our analyses and the number of observations per species are shown in Figure 3.

**Table 6.**
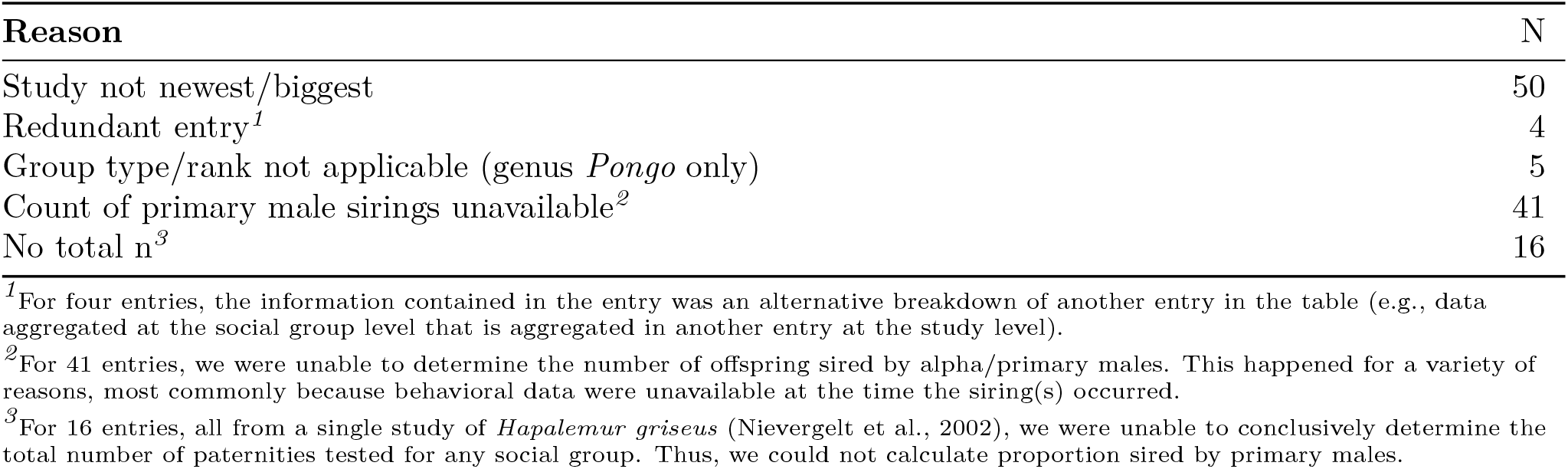
Reasons table entries were excluded from current analyses.

**Figure 3:**
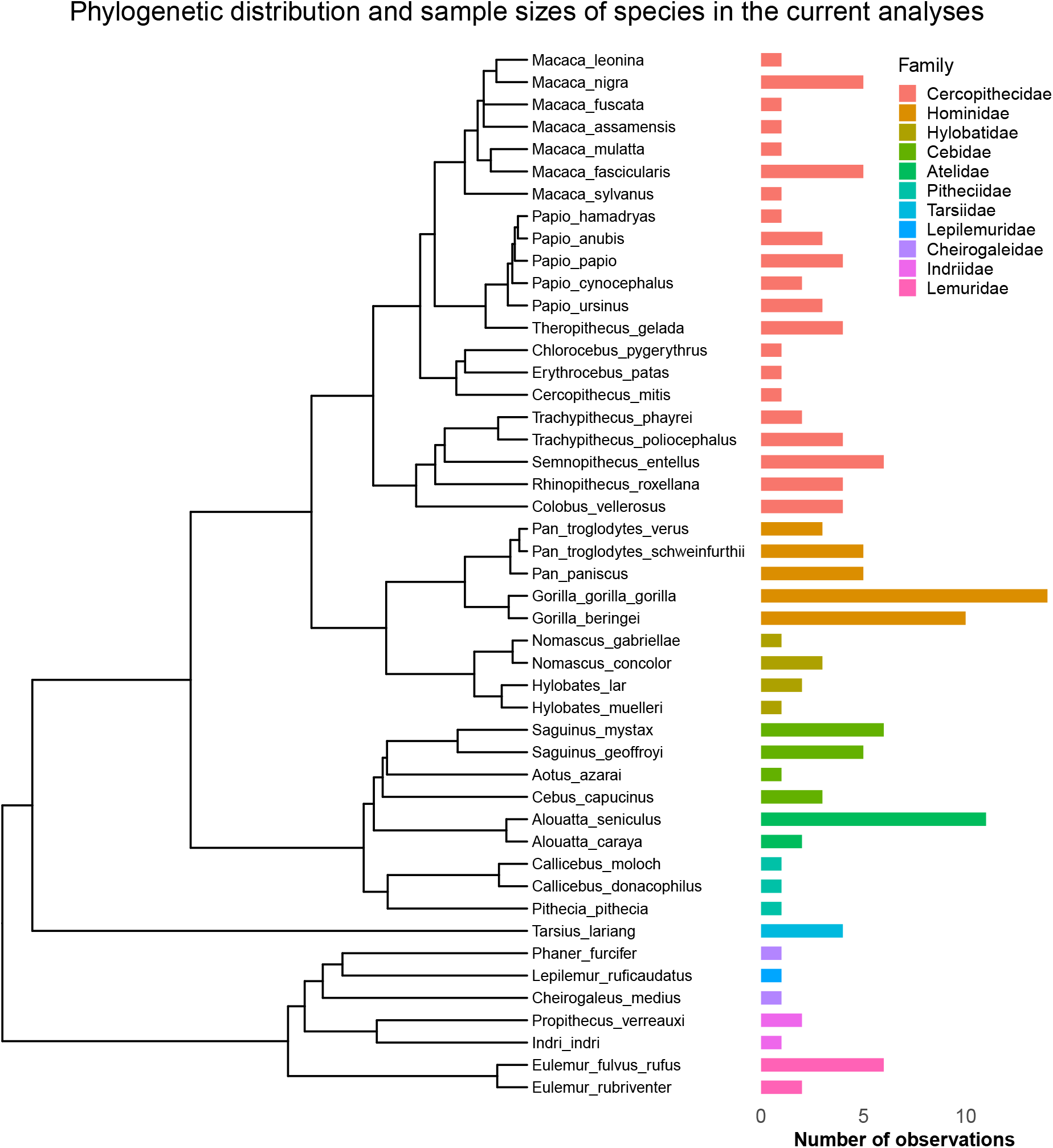
Phylogenetic distribution and sample sizes of species in our analyses. The left panel shows the consensus phylogeny from 10kTrees. The right panel shows the number of observations (rows in the dataset) per species, color-coded by family. The sample size in this figure does not include n=116 observations that were excluded from our analyses for the reasons listed in Table 6. Six species (toque macaque, gentle lemur, ring-tailed lemur, northern muriqui, and Bornean and Sumatran orangutans) included in the larger database are not represented in the present analyses.

### Question 1: To what degree does phylogeny predict the proportion of paternities obtained by primary males?

Species-level random effects, which include both phylogenetic relatedness and other species-level differences, explained roughly 35-40% of the variation in the proportion of paternities obtained by primary males. The posterior mean *R*^2^ values of intercept-only models for all group types when all uncertain paternities were assigned to the primary male (0.37, 95% CI: 0.31 - 0.43, n=148 observations, 46 species, and 2,432 total paternities) was very similar to the *R*^2^ value when none of the uncertain paternities were assigned to the primary male (0.39, 95% CI: 0.34 - 0.45).

We next examined how this explained variation was apportioned between phylogenetic relatedness and other, non-phylogenetic species-level factors (i.e., a higher *λ* represents a greater share of random effects explained by phylogeny). If we assume that primary males sired all infants of uncertain paternity, a substantial proportion of the variation in primary male paternity share was explained by phylogenetic relatedness in the full sample (*λ*=0.68, 95% CI: 0.18 - 0.93). Assigning all uncertain paternities to other, non-primary males decreased *λ* and increased 95% CIs (*λ*=0.52, 95% CI: 0.03 - 0.88). The remaining models presented in the results all use the same random effects structure that accounts for phylogenetic relatedness.

### Question 2: How do group composition and reproductive seasonality influence primary males’ paternity share?

#### Analysis 1: Effects of group composition on primary males’ paternity share

For this analysis, we grouped the multi-male/single-female observations (n=2) and multi-male/multi-female observations (n=81) into a single multi-male category. The multi-male/single-female groups could not be analyzed as a separate category due to the small sample size, and both observations had values that were well within the range for multi-male/multi-female groups (Table 7, Figure 4).

**Table 7.** Proportion of paternities obtained by primary males, by group type (raw data)

| Group composition | N obs | N species | Total paternities | Observed primary male paternity share |  |  |  |
| --- | --- | --- | --- | --- | --- | --- | --- |
|  |  |  |  | Min | Max | Mean (SD) min estimate | Mean (SD) max estimate |
| **Pair-living (all)** | 17 | 14 | 224 | 0.00 | 1.00 | 0.77 (0.28) | 0.77 (0.28) |
| cohesive | 10 | 10 | 157 | 0.74 | 1.00 | 0.91 (0.09) | 0.91 (0.09) |
| dispersed | 7 | 4 | 67 | 0.00 | 1.00 | 0.56 (0.33) | 0.56 (0.33) |
| **Single-male/multi-female** | 48 | 18 | 565 | 0.00 | 1.00 | 0.81 (0.26) | 0.82 (0.25) |
| **Multi-male/single-female** | 2 | 2 | 19 | 0.50 | 0.93 | 0.72 (0.31) | 0.72 (0.31) |
| **Multi-male/multi-female** | 81 | 26 | 1624 | 0.00 | 1.00 | 0.65 (0.27) | 0.71 (0.25) |

Note: Min and max are the lowest and highest observed primary male paternity shares across studies within each group type. Mean min estimate assigns no uncertain paternities to primary males; mean max estimate assigns all uncertain paternities to primary males.

**Figure 4:**
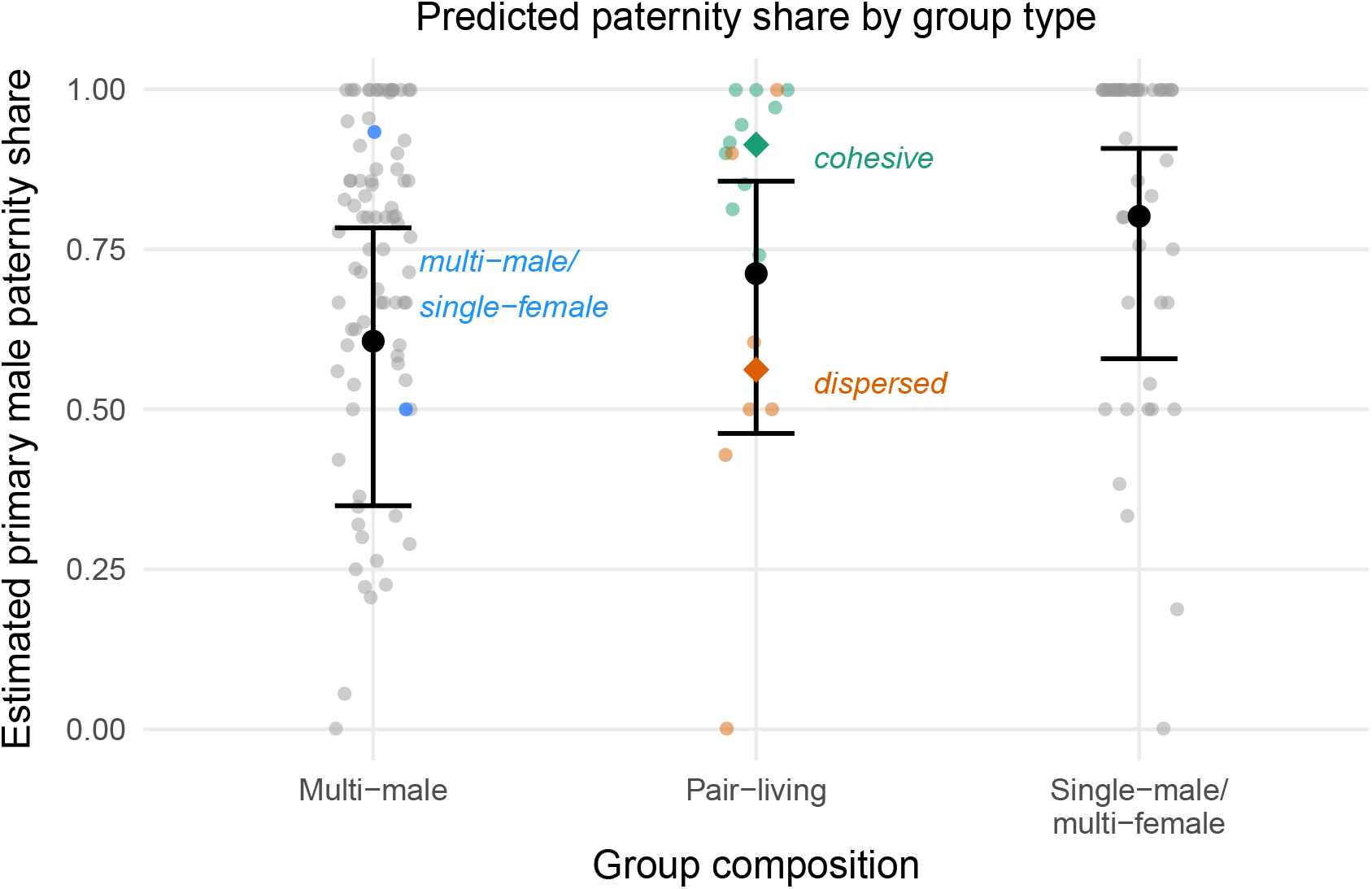
Males in multi-male groups had the lowest paternity share in the raw data (the small circles) and in model-derived predicted effects (black bars). The difference in paternity share between multi-male and single-female/multi-males groups was credible (i.e., the 95% CI did not include zero), while the credible intervals for the other two comparisons (multi-male to pair-living, and pair-living to single-male/multi-female) did not. The raw data mean for cohesive pair-living species is represented by a green diamond, while the mean for dispersed pair-living species is represented by an orange diamond.

In the raw data, primary males obtained on average 75% (SD = 26%) of paternities across all group types (Table 7); we obtained a similar value when all uncertain paternities were assigned to other males (mean = 72%, SD = 28%).

Primary males’ predicted paternity share varied with group composition. Primary males in single-male/multi-female groups had the highest average paternity share (posterior mean=0.79, 95% CI: 0.58 - 0.91) and those in multi-male groups the lowest average share (posterior mean=0.60, 95% CI: 0.35 - 0.78), though the variation within each category was considerable (Table 7, Figure 4). The difference between single-male/multi-female and multi-male groups was credible (95% CI: -0.28 - -0.11), while the evidence for a difference between pair-living and multi-male groups was weaker, with a 95% CI that barely crossed zero (-0.24 - 0.02). The relatively low paternity share in pair-living species was driven by dispersed pairs rather than cohesive pairs (Figure 4). See Tables S2-S5 in the supplementary materials for full results.

#### Analysis 2: Evaluation of the presence of in-group competitors and seasonality

We next examined the relationship between in-group competitors, seasonality, and primary male paternity share. Seasonality data were not available for all species, so this analysis was based on a slightly smaller sample than Analysis 1 (145 observations, 43 species, and 2,402 total paternities). In these models, both types of single-male groups (i.e., pair-living and single-male/multi-female) were combined into one category called single-male for two reasons: Analysis 1 estimates of primary males’ paternity shares in these groups were not reliably different, and they share the same key characteristic of having no in-group competitors.

The extent of reproductive seasonality had little effect on primary male paternity share, regardless of whether males lived in single or multi-male groups (Figure 5). Consistent with this overall conclusion, the estimated change in paternity share per unit increase in seasonality was near zero in both multi-male groups (posterior mean = 0.00, 95% CI: -0.16 - 0.15) and in single-male groups (posterior mean = 0.01, 95% CI: -0.13 - 0.14). The results were qualitatively very similar when pair-living groups were excluded, and when we used a more granular five-category version of the seasonality variable.

**Figure 5:**
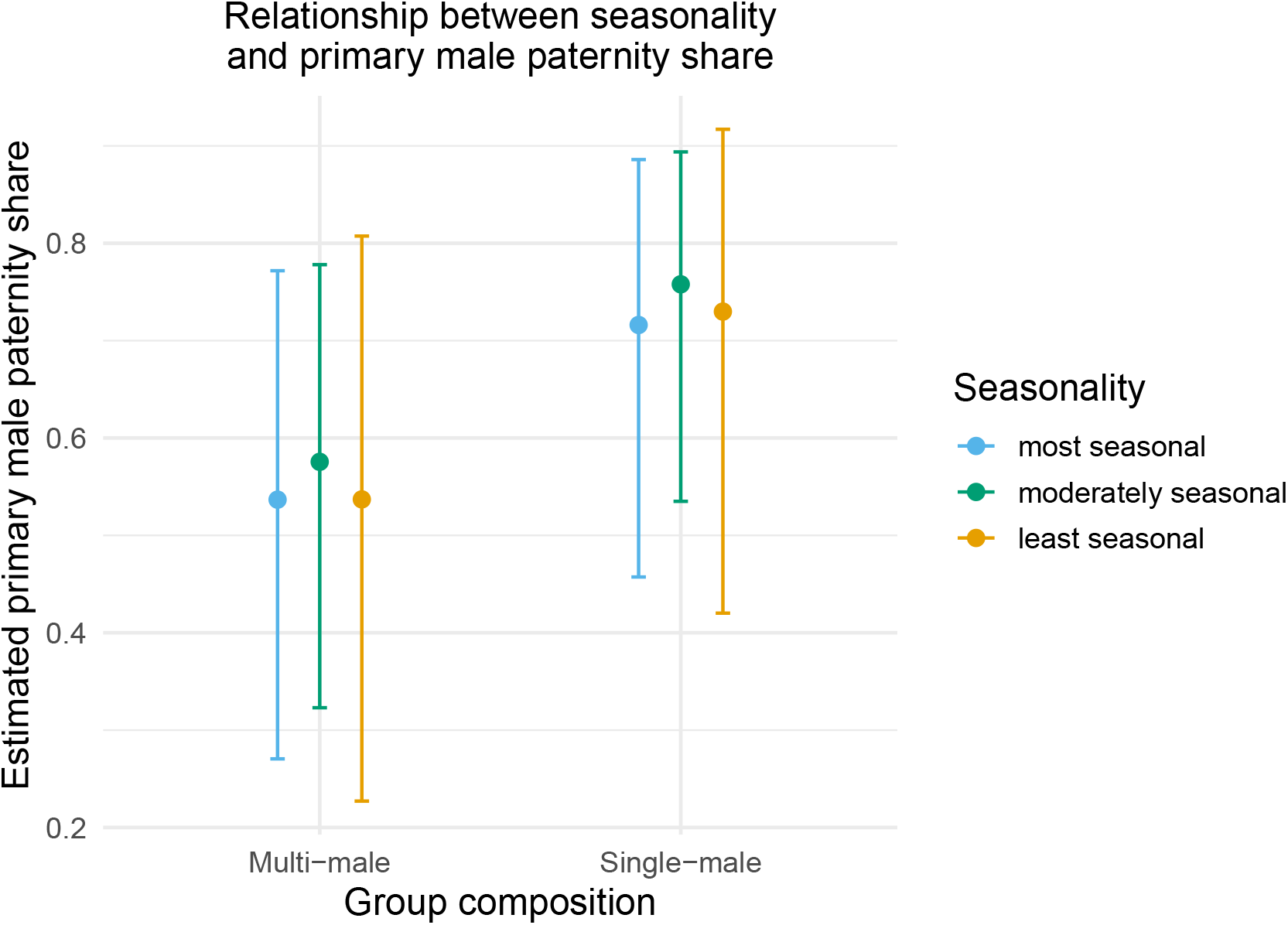
Seasonality had little effect on primary male paternity share. In both multi-male and single-male groups, species that were least seasonal, moderately seasonal, and most seasonal had similar estimated primary male paternity proportions, with extensive overlap in credible intervals. The results were qualitatively very similar when pair-living species were excluded from the single-male category.

The inclusion of seasonality in the model did not appreciably change the expected paternity share for primary males relative to what we found in Analysis 1 (Figure 5). Here, for multi-male groups the posterior mean was 0.58 (95% CI: 0.32 - 0.78) (versus 0.60 in Analysis 1). For single-male groups the posterior mean was 0.76 (95% CI: 0.53 - 0.89). This number is not directly comparable to a single Analysis 1 number because single-male/multi-female and pair-living observations are grouped together, but the mean is roughly halfway in between the Analysis 1 posterior mean values for pair-living males (0.70) and males in single-male/multi-female groups (0.79). The 95% CI of the difference between multi-male and single-male groups did not include zero (95% CI: 0.10 - 0.27), indicating a credible difference. See Tables S6-S9 in the supplementary materials for full results.

### Question 3: When primary males in multi-male groups lose paternities, to whom are they losing them?

In the raw data, a minimum of 6% (SD = 17%) of paternities in multi-male groups were assigned to extra-group males; this rose to 9% (SD = 20%) if all uncertain paternities were assigned to extra-group males (n=90 observations, 31 species, and 1,758 total paternities). Primary males in multi-male groups sire approximately 60% of infants (i.e., the posterior mean prediction from Question 2, Analysis 1 above), so about 40% of infants are sired by other males. If the raw EGP rate is 6–9%, this would mean that 15–22% of the paternities that primary males lose are attributable to extra-group males, with the remainder lost to group mates.^2^

While our statistical models generally tell a similar story, they suggest that losses to extra-group males may be slightly higher than values obtained from the raw data. For the subset of data where all relevant information was available (66 observations, 27 species, and 1,419 paternities), an estimated 26% of primary males’ lost paternities were EGPs (95% CI: 4 - 69%). The other 74% of lost paternities went to in-group males (95% CI: 31 - 96%).

Since the 95% CI for primary male paternity share in multi-male groups is 35-78% (see Figure 4), this means that somewhere between 22% and 65% of infants are sired by other males. If 26% of the low end and high end of these paternity share estimates are EGPs, this would translate to an overall EGP rate of ∼6-17%.^3^ See Tables S10-S13 in the supplementary materials for full results.

## Discussion

The primary goal of this project was to compile all available genetic paternity data for wild nonhuman primates into a single publicly accessible repository. The database that we have created contains information about 52 species from 31 genera representing 11 of 14 primate families in 10kTrees. One core feature of our dataset is that it acknowledges the ambiguity and uncertainty in the data that arises when not all sires are identified, not all potential sires can be excluded, dominance ranks at the time of conception are not known (or reported), and not all paternities are resolved with high degrees of certainty. This enables researchers to bracket estimates of paternity metrics between plausible minimum and maximum values. In addition, we tabulated data at the group level whenever possible. This provides information about magnitude of the range of variation within and between groups, populations, and species.

Despite the large number of species in our dataset, we are missing information from a substantial number of taxa. To the best of our knowledge, no data are available for any species in three primate families (*Galagidae, Lorisidae*, and *Daubentoniidae*). While most species in these families are solitary and thus not relevant for assessing primary males’ paternity share or the rate of extra-group paternities, in some a single male might control a territory that encompasses the ranges of multiple females (like orangutans; see e.g., Noordwijk et al. (2023)). We are also missing data from many species of other primate taxa that live in social groups, and some families, particularly the *Cercopithecidae* and *Hominidae*, are over-represented in the data.

Phylogeny has a detectable but moderate effect on the distribution of paternity in primate groups. Species-level random effects, which do not differentiate between phylogenetic and non-phylogenetic species differences, explained roughly 35–40% of the total variation in primary males’ paternity share. Our models indicate that between 52% and 68% of that inter-species variation is attributable to phylogenetic relationships. In absolute terms, this means phylogeny accounts for a relatively modest share of total variation in paternity outcomes. Instead, most of the variation comes from observation-level differences within and between species that are not captured by the phylogenetic or species-level random effects. Kamilar and Cooper (2013) found a strong phylogenetic signal associated with morphological traits related to the intensity of intrasexual selection on males, including sexual dimorphism in body mass (*λ* = 0.94), canine size (*λ* = 0.82), and testes size (*λ* = 0.89). They also reported a Pagel’s *λ* of 0.60 for the number of males in groups. Because our *λ* is computed differently than Pagel’s *λ* (i.e., on the random effects derived from our statistical model, rather than directly from the phylogenetic tree), we can’t directly compare our estimates to those values. However, the general pattern is consistent: morphological traits associated with male-male competition show very strong phylogenetic signal, while behavioral and demographic traits like group composition and paternity share show comparatively weaker signals.

### Primary males’ paternity share is strongly influenced by the kinds of groups in which they live

Interestingly, pair-living non-human primate species encompass both lowest and highest rates of paternity loss for primary males. For species that live in cohesive pairs, primary males sire about 90% of their partners’ offspring. For species that live in dispersed pairs, primary males sire only about 55%. Similar findings have been reported for a broader range of pair-living mammalian species (Clutton-Brock and Isvaran 2006; Huck et al. 2014). Cohas and Allainé (2009) found no difference between pair-living species that form dispersed and cohesive pair bonds, but they reported that solitary species have much higher rates of extra-group paternities than pair-living species. However, they categorized as solitary at least one species, the fork-marked lemur (*Phaner furcifer*), that other researchers have described as having dispersed pair bonds (Schülke and Kappeler 2003). Thus, the preponderance of evidence suggests that for pair-living mammals, paternity patterns are strongly influenced by association patterns. This reinforces the need to recognize that pair-living (or social monogamy) is not a homogeneous phenomenon, a point which has been made before by others (e.g., Kappeler 2014; Fernandez-Duque et al. 2020; Huck, Di Fiore, and Fernandez-Duque 2020).

Males in single-male/multi-female groups sire the great majority of offspring within their groups, but do not monopolize paternity as successfully as males in cohesive pairs do. This is likely to reflect the fact that competition over access to groups of females is more intense than competition over access to single females and it is harder for males to monopolize multiple females than to monopolize single females. Blue monkeys, for example, typically form single-male/multi-female groups, but there are regular incursions by outside males during the mating season. Larger numbers of sexually receptive females draw larger numbers of outside males (Gao and Cords 2020).

Primary males living in multi-male groups lose more paternities than primary males living in cohesive pairs or in single-male/multi-female groups, similar to patterns that have been reported for mammals more generally (Clutton-Brock and Isvaran 2006). These losses may be the product of several different mechanisms. First, primary males may not be able to monopolize paternity when more than one sexually receptive female is present (Altmann 1962). Overlap in female receptivity is particularly likely to occur in multi-male/multi-female groups because there are larger numbers of females in these groups (Ostner, Nunn, and Schülke 2008). Second, in species with long male tenures, females may avoid mating with closely related males. The tenures of primary males in white faced capuchin groups are sometimes long enough that females live in the same groups as their fathers, and males don’t sire the offspring of closely related females (Godoy, Vigilant, and Perry 2016; Wikberg et al. 2017). Third, males in multi-male groups may form coalitions that influence their access to sexually receptive females. Middle-ranking male olive and yellow baboons sometimes team up to disrupt consortships of high-ranking males (Packer 1977; Noë 1992), although we don’t know whether these consort takeovers influence paternity. Male chimpanzees rely on their social ties with other males to obtain and maintain high rank and to defend the boundaries of their territories. Alpha males selectively tolerate matings by their allies (Duffy, Wrangham, and Silk 2007; Bray, Pusey, and Gilby 2016), and this translates into paternity success for males with close ties to the alpha male (Feldblum et al. 2021).

### Primary males in multi-male groups mostly lose paternities to other resident males

Our data also show that primary males in multi-male groups lose the majority of paternities to other males within their groups, not to extra-group males. This may be because resident males actively deter incursions by outside males or because subordinate residents are better able to capitalize on reproductive opportunities than outside males. For example, male chacma baboons “monitor consortships so assiduously that they rapidly recognize both when an unexpected mating opportunity arises and when a consortship has ended” (Crockford et al. 2007, 889). Their attentiveness may give them an important advantage over extra-group males.

### Seasonality has little effect on primary males’ paternity share

Contrary to our predictions, the extent of reproductive seasonality had little effect on the paternity share of primary males. This prediction was based on the assumption that when reproductive seasonality was strong, there would be greater overlap in females’ sexual receptivity, and this would limit primary males’ ability to monopolize conceptions. In retrospect, there are reasons to be skeptical of this assumption. Though at least one mammalian meta-analysis found evidence that EGPs increase when mating seasons are shorter (Isvaran and Clutton-Brock 2007), primate-specific studies have provided mixed evidence about the effects of reproductive seasonality on males’ ability to monopolize access to females (Paul 1997; Kutsukake and Nunn 2006; Ostner, Nunn, and Schülke 2008; Gogarten and Koenig 2013). Gogarten and Koenig (2013) showed that the extent of reproductive synchrony among females does not vary consistently with the extent of reproductive seasonality.

It seems counter-intuitive that reproductive seasonality is unrelated to the extent of reproductive synchrony among females. If conceptions are more concentrated in time, then there should be more overlap in reproductive synchrony among females. However, the extent of reproductive synchrony will be a joint function of length of the breeding season, the duration of sexual receptivity or the conception window, the number of females in groups, and males’ ability to distinguish between conceptive and nonconceptive cycles and to identify conception windows. Thus, a highly seasonal species with very short conception windows, small group size, and male knowledge of conception timing might have relatively little effective reproductive synchrony. At the same time, a species with little reproductive seasonality might have long periods of sexual receptivity and effective concealment of the timing of conception, which would generate considerable reproductive synchrony among females.

To assess the impact of reproductive synchrony on primary males’ ability to monopolize conceptions, we need more detailed information about the number of sexually receptive or conceiving females that are present in groups on days in which there is at least one sexually receptive or conceiving female present, as well as males’ ability to identify the timing of conceptions. These kinds of data are not widely available.

### Limitations of the study

Our analysis has a number of limitations. First, almost all of the samples are incomplete in some way. In some cases, not all infants born in the population during the study period were sampled, not all potential sires were sampled, or it was not possible to exclude more than one male as the sire of a particular infant. Second, most species that are included in the dataset are represented by a single study population, and sometimes a single group. This may bias estimates of the range of variation within and between species (see Alberts, Watts, and Altmann 2003). Third, many species are missing from our database and certain taxa are over-represented.

In addition, the exercise of writing out our theoretical model made clear the significant disconnect between the data that would ideally be available to answer our questions and the data that are actually available. An ideal dataset would document for each infant the date of conception, the rank and identity of the sire, the certainty of the paternity estimate, the number of potential sires in the group at the time of conception, and the number of sexually receptive females present in the group at the time of conception. We welcome the opportunity to incorporate such data into our dataset.

## Conclusions

We have constructed the most complete dataset of genetic paternity data for wild nonhuman primates to date. As a “living” and publicly accessible resource, the database provides a template for researchers to contribute new paternity information and to fill gaps in previously published paternity data.

Our analyses show that both phylogeny and group composition influence the paternity share of primary males. The most effective tactic for monopolizing conceptions is to live in cohesive pairs and the least effective tactic is to live in groups with other males. As expected based on prior work, rates of extra-group paternities are lower in multi-male groups than in other kinds of groups. However, males’ options are constrained by ecological factors which influence the distribution of sexually receptive females in space and time. As more complete information about genetic paternity becomes available for more groups and species, we will be able to construct more powerful analyses of the forces that shape male reproductive strategies.

## Acknowledgements

We would like to thank Adrian Jäggi, Eduardo Fernandez-Duque, and Jason Kamilar for their advice and thoughts on these analyses, as well as Stephanie Fox and Pascale Sicotte, Carola Borries, Sylvain Gatti and colleagues, Liza Moscovice and Robert Seyfarth, Veronika Städele, and Linda Vigilant for contributing unpublished paternity data. Maud Mouginot, Amy Scott, Joseph Feldblum, Songtao Guo, Eva Wikberg, Peter Kappeler, Caroline Nievergelt, and Carola Borries all helped us untangle and extract data from published papers, for which we are extremely grateful.

## Data and code availability

All data, code, and supplementary materials are available at https://github.com/slrosen/primate-paternity-metanalysis.

## GenAI use statement

We used Claude Code to help streamline our data cleaning and compilation workflow, clean and edit code, debug output formatting issues, alphabetize and consolidate/organize .bib files and the associated study pdfs, copy edit the text, and create and update the README file that is available on Github. Our CLAUDE.md file can be found on Github along with all other materials needed to replicate these analyses.

## Footnotes

1 Occasionally the names were different, for any of three reasons: 1) 10kTrees used an older classification scheme (e.g., the older, monogeneric *Callicebus* classification for titi monkeys, while the study we included uses *Plecturocebus* (Dolotovskaya, Roos, and Heymann 2020)); 2) 10kTrees did not contain an exact match (e.g., it did not have *Brachyteles hypoxanthus*, so we used *B. arachnoides*); or 3) the study omitted subspecies name while 10kTrees required it (e.g., few published studies distinguish between *Pan troglodytes* subspecies, while 10kTrees requires specification of either P. t. verus or *P. t. schweinfurthii*).

2 6/40 = 15%; 9/40 = 22%.

3 0.26 × 0.22 = 0.06; 0.26 × 0.65 = 0.17.

