## Supplementary materials for "Distribution of genetic paternity in primate groups"

Stacy Rosenbaum, Nicholas Grebe, and Joan B. Silk

**The main text for these supplementary materials can be accessed at https://github.com/slrosen/primate-paternity-metanalysis.**

Document rendered on April 2, 2026

### 1. Bayesian model diagnostics

Table S1 summarizes convergence diagnostics for the Bayesian models reported in the main text. Q1 refers to the models used to answer Question 1 in the results in the main text; Q2/A1 and Q2/A2 refer to Question 2, Analysis 1 and 2 respectively, and Q3 refers to Question 3. Figures S1–S5 show the posterior predictive checks for each of these models. In each plot, the dark blue line represents the observed data distribution, and the lighter blue lines represent distributions simulated from the fitted model's posterior. Significant overlap between the observed and simulated distributions indicates that the model captured the important features of the data.

Table S1. Bayesian model convergence diagnostics

| Model | Max Rhat | Min Bulk ESS | Min Tail ESS | Divergent transitions |
|---|---|---|---|---|
| Q1: Phylogeny (Intercept-only, max) | 1.0011 | 3414 | 5076 | 0 |
| Q1: Phylogeny (Intercept-only, min) | 1.0029 | 2341 | 3157 | 0 |
| Q2/A1: Group composition | 1.0003 | 3915 | 4164 | 0 |
| Q2/A2: Competition/seasonality | 1.0004 | 4176 | 4588 | 0 |
| Q3: Paternity loss source | 1.0002 | 4187 | 4666 | 0 |

##### Posterior Predictive Checks

Figure S1. Q1 Phylogeny (Intercept−only, max)

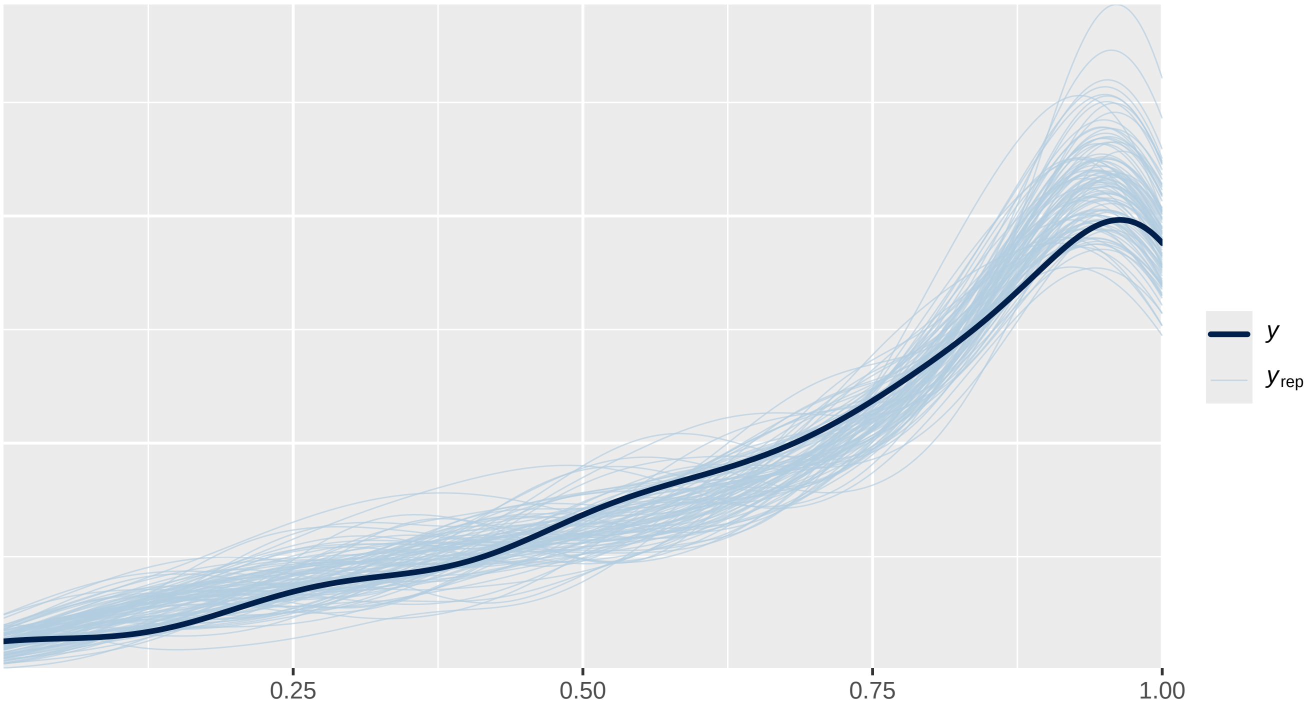

Figure S2. Q1 Phylogeny (Intercept−only, min)

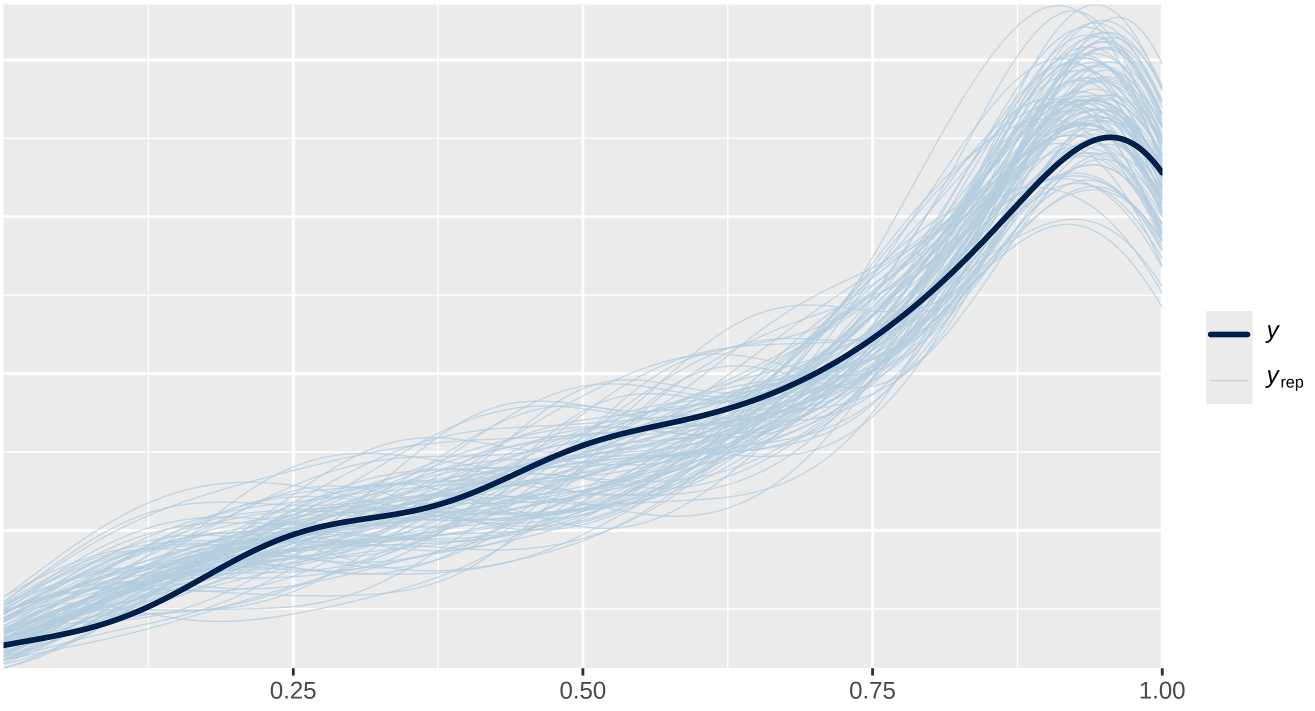

Figure S3. Q2/A1 Group composition

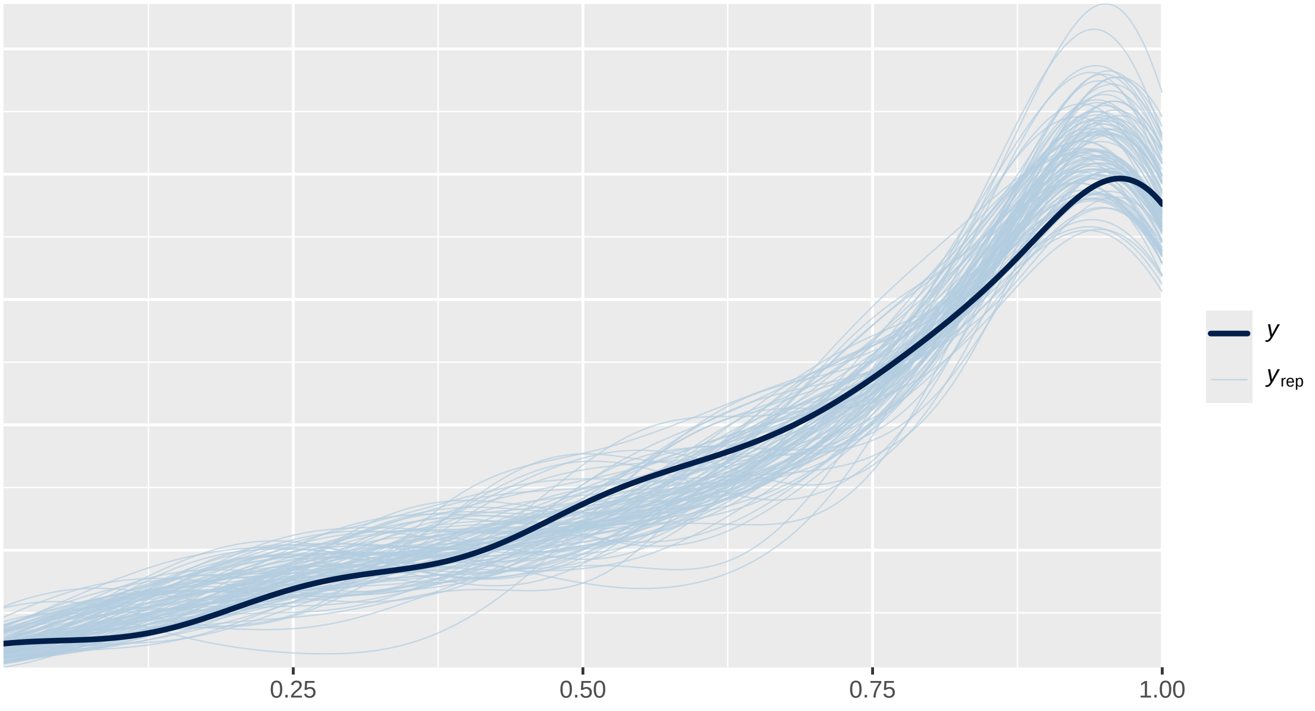

Figure S4. Q2/A2 Competition/seasonality

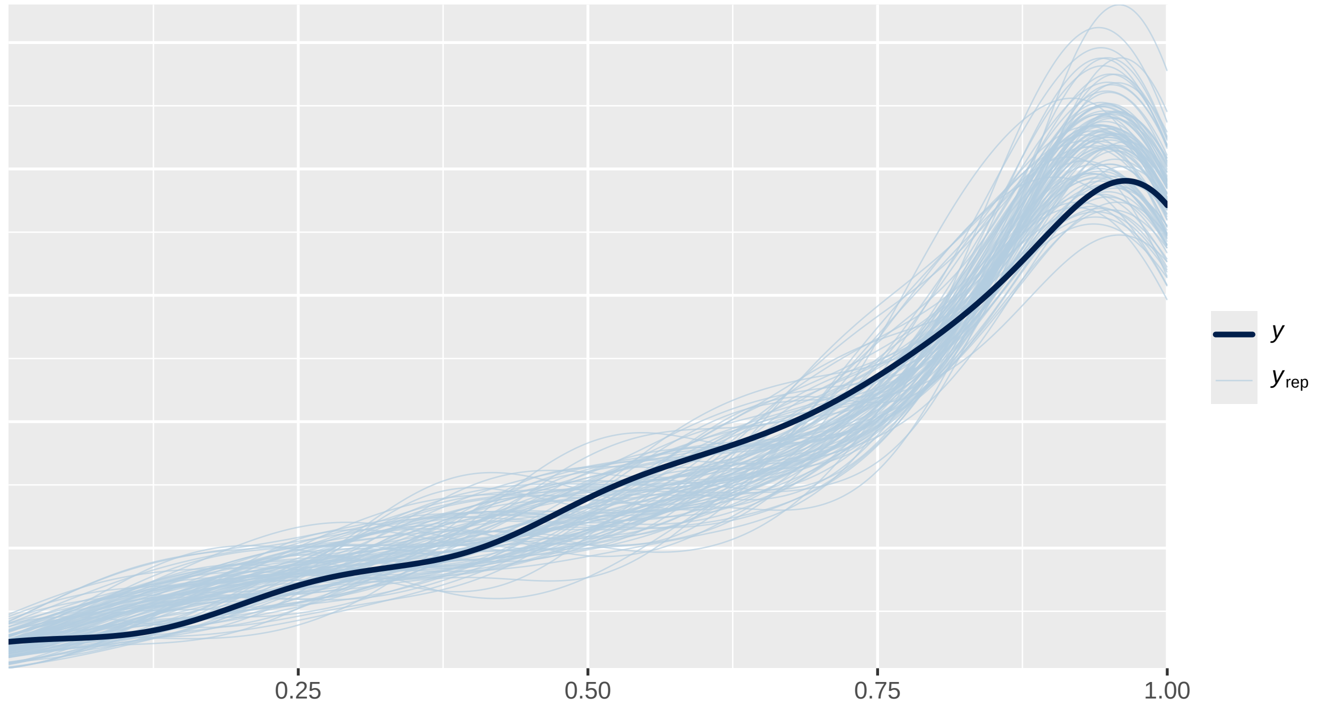

Figure S5. Q3 Paternity loss source

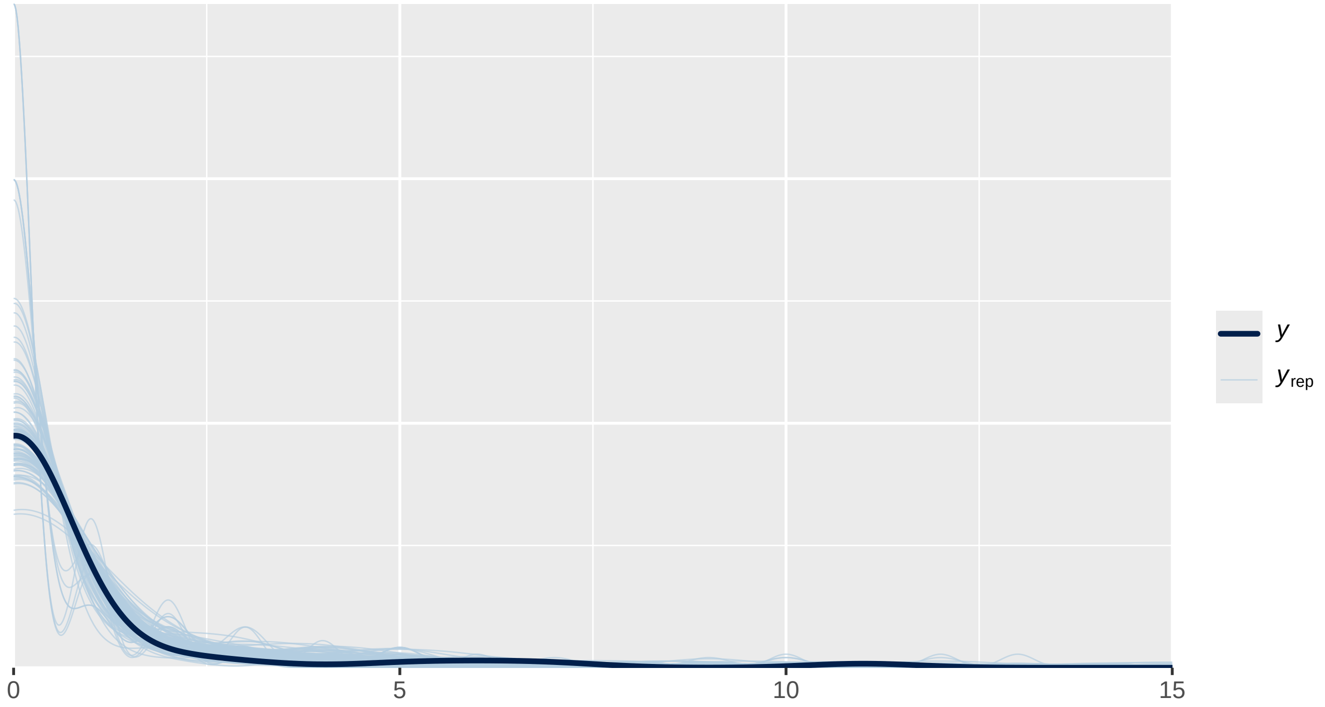

### 2. Sensitivity analyses: assignment of uncertain paternities

#### A. The effects of group composition (Question 2, Analysis 1 in main text)

In the main text the models for Question 2 use the maximum possible estimate of primary male paternity share; i.e., all uncertain paternities are assigned to primary males. For comparison purposes, here we present results from those models alongside results from models using minimum estimates, in which none of the uncertain paternities are assigned to primary males.

Table S2. Model results: Maximum primary male paternity share (in main text)

| Parameter | Estimate | Est.Error | l-95% CI | u-95% CI | Rhat | Bulk_ESS | Tail_ESS |
|---|---|---|---|---|---|---|---|
| Intercept (Multi-male) | 0.405 | 0.476 | -0.622 | 1.286 | 1.000 | 17,269 | 13,670 |
| Pair-living | 0.482 | 0.300 | -0.096 | 1.074 | 1.000 | 26,371 | 15,790 |
| Single-male/multi-female | 0.964 | 0.172 | 0.628 | 1.307 | 1.000 | 29,322 | 14,819 |

Table S3. Model results: minimum primary male paternity share

| Parameter | Estimate | Est.Error | l-95% CI | u-95% CI | Rhat | Bulk_ESS | Tail_ESS |
|---|---|---|---|---|---|---|---|
| Intercept (Multi-male) | 0.244 | 0.434 | -0.723 | 1.032 | 1.001 | 12,258 | 10,985 |
| Pair-living | 0.705 | 0.291 | 0.136 | 1.277 | 1.000 | 19,555 | 15,331 |
| Single-male/multi-female | 1.333 | 0.165 | 1.010 | 1.660 | 1.000 | 23,841 | 15,093 |

Table S4. Model-derived predicted paternity share using maximum estimate (in main text)

| Group type | Mean | 95% CI |
|---|---|---|
| Multi-male | 59.6% | 34.9% - 78.3% |
| Pair-living | 69.9% | 46.2% - 85.6% |
| Single-male/multi-female | 78.6% | 57.9% - 90.8% |

Table S5. Model-derived predicted paternity share using minimum estimate

| Group type | Mean | 95% CI |
|---|---|---|
| Multi-male | 55.9% | 32.7% - 73.7% |
| Pair-living | 71.3% | 49.2% - 85.4% |
| Single-male/multi-female | 82.0% | 63.9% - 91.6% |

#### B. The effects of in-group competition and seasonality (Question 2, Analysis 2 in main text)

Table S6. Model results: Maximum primary male paternity share (in main text)

| Parameter | Estimate | Est.Error | l-95% CI | u-95% CI | Rhat | Bulk_ESS | Tail_ESS |
|---|---|---|---|---|---|---|---|
| Intercept | -0.332 | 0.958 | -2.214 | 1.542 | 1.000 | 23,627 | 15,987 |
| Competition[1] | 0.780 | 0.479 | -0.152 | 1.702 | 1.000 | 21,751 | 14,993 |
| Seasonality | 0.649 | 0.824 | -0.968 | 2.249 | 1.000 | 20,675 | 15,702 |
| Seasonality^2 | -0.162 | 0.214 | -0.578 | 0.263 | 1.000 | 18,262 | 15,203 |
| Competition*Seasonality | 0.051 | 0.239 | -0.415 | 0.524 | 1.000 | 21,846 | 14,552 |

[1]Single-male versus multi-male group

Table S7. Model results: minimum primary male paternity share

| Parameter | Estimate | Est.Error | l-95% CI | u-95% CI | Rhat | Bulk_ESS | Tail_ESS |
|---|---|---|---|---|---|---|---|
| Intercept | -0.208 | 0.953 | -2.075 | 1.687 | 1.000 | 16,474 | 14,827 |
| Competition[1] | 0.908 | 0.458 | 0.014 | 1.809 | 1.000 | 15,966 | 14,648 |
| Seasonality | 0.493 | 0.809 | -1.118 | 2.068 | 1.000 | 14,675 | 14,878 |
| Seasonality^2 | -0.177 | 0.204 | -0.576 | 0.229 | 1.000 | 13,414 | 13,925 |
| Competition*Seasonality | 0.169 | 0.230 | -0.281 | 0.617 | 1.000 | 15,660 | 14,594 |

[1]Single-male versus multi-male group

**Comparing the conditional effect of in-group competition between models** Because the competition/seasonality models presented above contain interaction terms, the main effect of competition cannot be interpreted in isolation. In the below tables we compute the conditional effect of competition (i.e., the difference in predicted paternity share between single-male and multi-male groups) at different levels of seasonality for both the maximum and minimum paternity share models. The purpose of this is to determine whether the difference in the main effect coefficient on the competition parameter leads to a substantively different conclusion.

Table S8. Conditional effect of competition (maximum estimate)

| Seasonality | Multi-male predicted | Single-male predicted | Difference | 95% CI |
|---|---|---|---|---|
| Most seasonal | 53.7% | 71.6% | 17.9% | 6.2% to 30.3% |
| Moderately seasonal | 57.6% | 75.8% | 18.2% | 9.8% to 27.0% |
| Least seasonal | 53.7% | 73.0% | 19.3% | 6.1% to 33.9% |

Table S9. Conditional effect of competition (minimum estimate)

| Seasonality | Multi-male predicted | Single-male predicted | Difference | 95% CI |
|---|---|---|---|---|
| Most seasonal | 52.5% | 75.3% | 22.8% | 10.9% to 34.8% |
| Moderately seasonal | 51.7% | 77.8% | 26.1% | 16.7% to 35.0% |
| Least seasonal | 42.7% | 73.1% | 30.4% | 16.1% to 44.0% |

**Interpretation** In both models, competition has a meaningful effect on primary male paternity share: males in single-male groups are predicted to obtain a substantially higher share of paternities than males in multi-male groups at all levels of seasonality. However, the minimum estimate model produces larger point estimates for this effect (~23–30% difference versus ~18–19% difference in the maximum estimate model). This is not surprising. Reassigning uncertain paternities away from primary males disproportionately reduces the predicted primary male paternity share in multi-male groups, where most of the ambiguity in paternity assignment occurs. However, there is considerable overlap in the 95% credible intervals between the two models at every level of seasonality. Thus, these two different methods of operationalizing primary male paternity share do not ultimately yield distinguishable estimates of the effect of competition. The qualitative conclusion (that competition is an important predictor of primary male paternity share, while seasonality is not) is robust to decisions about how to handle uncertain paternities.

#### C. Source of paternity losses in multi-male groups (Question 3 in main text)

In the main text we model the proportion of lost paternities attributable to extra-group males versus non-primary resident males using the minimum extra-group paternity (EGP) count (i.e., only confirmed extra-group paternities). Here we present that model alongside a model using the maximum possible EGP count, in which ambiguous paternities are assigned to extra-group males. Results are qualitatively similar regardless of which choice is made.

Table S10. Paternity loss source model: minimum EGP count (in main text)

| Parameter | Estimate | Est.Error | l-95% CI | u-95% CI | Rhat | Bulk_ESS | Tail_ESS |
|---|---|---|---|---|---|---|---|
| Intercept | -1.294 | 1.060 | -3.290 | 0.823 | 1.000 | 9,531 | 10,218 |

Table S11. Paternity loss source model: maximum EGP count

| Parameter | Estimate | Est.Error | l-95% CI | u-95% CI | Rhat | Bulk_ESS | Tail_ESS |
|---|---|---|---|---|---|---|---|
| Intercept | -1.055 | 0.889 | -2.620 | 0.818 | 1.000 | 6,798 | 12,667 |

Table S12. Model-derived predicted paternity loss source: minimum EGP count (in main text)

| Loss source | Mean | 95% CI |
|---|---|---|
| Extra-group males | 25.7% | 3.6% - 69.5% |
| Non-primary resident males | 74.3% | 30.5% - 96.4% |

Table S13. Model-derived predicted paternity loss source: maximum EGP count

| Loss source | Mean | 95% CI |
|---|---|---|
| Extra-group males | 28.6% | 6.8% - 69.4% |
| Non-primary resident males | 71.4% | 30.6% - 93.2% |

### 3. Paternity data bibliography

The following references are the sources from which genetic paternity data were compiled for this study.

Alberts, Susan C., Jason C. Buchan, and Jeanne Altmann. 2006. "Sexual Selection in Wild Baboons: From Mating Opportunities to Paternity Success." *Animal Behaviour* 72: 1177–96. https://doi.org/10.1016/j.anbehav.2006.05.001.

Altmann, Jeanne, Susan C. Alberts, Susan A. Haines, Jean Dubach, Philip Muruthi, Trevor Coote, Eli Geffen, et al. 1996. "Behavior Predicts Genetic Structure in a Wild Primate Group." *Proceedings of the National Academy of Sciences of the United States of America* 93: 5797–5801. https://doi.org/10.1073/pnas.93.12.5797.

Banes, Graham L., Biruté M. F. Galdikas, and Linda Vigilant. 2015. "Male Orang-Utan Bimaturism and Reproductive Success at Camp Leakey in Tanjung Puting National Park, Indonesia." *Behavioral Ecology and Sociobiology* 69: 1785–94. https://doi.org/10.1007/s00265-015-1991-0.

Barelli, Claudia, Kazunari Matsudaira, Tanja Wolf, Christian Roos, Michael Heistermann, Keith Hodges, Takafumi Ishida, Suchinda Malaivijitnond, and Ulrich H. Reichard. 2013. "Extra-Pair Paternity Confirmed in Wild White-Handed Gibbons." *American Journal of Primatology* 75: 1185–95. https://doi.org/10.1002/ajp.22180.

Boesch, Christophe, Grégoire Kohou, Honora Néné, and Linda Vigilant. 2006. "Male Competition and Paternity in Wild Chimpanzees of the Taï Forest." *American Journal of Physical Anthropology* 130: 103–15. https://doi.org/10.1002/ajpa.20341.

Bonadonna, Giovanna, Valeria Torti, Chiara De Gregorio, Daria Valente, Rose Marie Randrianarison, Luca Pozzi, Marco Gamba, and Cristina Giacoma. 2019. "Evidence of Genetic Monogamy in the Lemur Indri (Indri Indri)." *American Journal of Primatology.* https://doi.org/10.1002/ajp.22993.

Borries, Carola. n.d. "Unpublished Paternity Data for Phayre's Leaf Monkeys."

Bradley, Brenda J., Diane M. Doran-Sheehy, Dieter Lukas, Christophe Boesch, and Linda Vigilant. 2004. "Dispersed Male Networks in Western Gorillas." *Current Biology* 14: 510–13. https://doi.org/10.1016/j.cub.2004.02.062.

Bradley, Brenda J., Martha M. Robbins, Elizabeth A. Williamson, H. Dieter Steklis, Netzin Gerald Steklis, Nadin Eckhardt, Christophe Boesch, and Linda Vigilant. 2005. "Mountain Gorilla Tug-of-War: Silverbacks Have Limited Control over Reproduction in Multimale Groups." *Proceedings of the National Academy of Sciences of the United States of America* 101: 9187–91. https://doi.org/10.1073/pnas.0502019102.

Brauch, Katrin, Keith Hodges, Antje Engelhardt, Kerstin Fuhrmann, Eric Shaw, and Michael Heistermann. 2008. "Sex-Specific Reproductive Behaviours and Paternity in Free-Ranging Barbary Macaques (Macaca Sylvanus)." *Behavioral Ecology and Sociobiology* 62: 1453–66. https://doi.org/10.1007/s00265-008-0575-7.

Cai, YanSen, HaoYang Yu, Hua Liu, Cong Jiang, Ling Sun, LiLi Niu, XuanZhen Liu, DaYong Li, and Jing Li. 2020. "Genome-Wide Screening of Microsatellites in Golden Snub-Nosed Monkey (Rhinopithecus Roxellana), for the Development of a Standardized Genetic Marker System." *Scientific Reports* 10. https://doi.org/10.1038/s41598-020-67451-2.

Chaves, P. B., K. B. Strier, and A. Di Fiore. 2023. "Paternity Data Reveal High MHC Diversity Among Sires in a Polygynandrous, Egalitarian Primate." *Proceedings of the Royal Society B* 290: 20231035. https://doi.org/10.1098/rspb.2023.1035.

Constable, Julie L., Mary V. Ashley, Jane Goodall, and Anne E. Pusey. 2001. "Noninvasive Paternity Assignment in Gombe Chimpanzees." *Molecular Ecology* 10: 1279–1300. https://doi.org/10.1046/j.1365-294X.2001.01262.x.

Dal Pesco, Federica. 2019. "Dynamics and Fitness Benefits of Male-Male Sociality in Wild Guinea Baboons (Papio Papio)." PhD thesis, Georg-August-Universität Göttingen.

Dal Pesco, Federica, Christof Neumann, Franziska Trede, Dietmar Zinner, and Julia Fischer. 2025. "Nested Male Reproductive Strategies in a Tolerant Multilevel Primate Society." *bioRxiv.* https://doi.org/10.1101/2025.10.27.684814.

Díaz-Muñoz, Samuel L. 2011. "Paternity and Relatedness in a Polyandrous Nonhuman Primate: Testing Adaptive Hypotheses of Male Reproductive Cooperation." *Animal Behaviour* 82: 563–71. https://doi.or

g/10.1016/j.anbehav.2011.06.013.

Dolotovskaya, Sofya, Christian Roos, and Eckhard W. Heymann. 2020. "Genetic Monogamy and Mate Choice in a Pair-Living Primate." *Scientific Reports* 10. https://doi.org/10.1038/s41598-020-77132-9.

Driller, Christine, Dyah Perwitasari-Farajallah, Hans Zischler, and Stefan Merker. 2009. "The Social System of Lariang Tarsiers (Tarsius Lariang) as Revealed by Genetic Analyses." *International Journal of Primatology* 30: 267–81. https://doi.org/10.1007/s10764-009-9341-6.

Engelhardt, Antje, Michael Heistermann, J. Keith Hodges, Peter Nürnberg, and Carsten Niemitz. 2006. "Determinants of Male Reproductive Success in Wild Long-Tailed Macaques (Macaca Fascicularis)–Male Monopolisation, Female Mate Choice or Post-Copulatory Mechanisms?" *Behavioral Ecology and Sociobiology* 59: 740–52. https://doi.org/10.1007/s00265-005-0104-x.

Engelhardt, Antje, Laura Muniz, Dyah Perwitasari-Farajallah, and Anja Widdig. 2017. "Highly Polymorphic Microsatellite Markers for the Assessment of Male Reproductive Skew and Genetic Variation in Critically Endangered Crested Macaques (Macaca Nigra)." *International Journal of Primatology* 38: 672–91. https://doi.org/10.1007/s10764-017-9973-x.

Feldblum, Joseph T., Emily E. Wroblewski, Rebecca S. Rudicell, Beatrice H. Hahn, Thais Paiva, Mine Cetinkaya-Rundel, Anne E. Pusey, and Ian C. Gilby. 2014. "Sexually Coercive Male Chimpanzees Sire More Offspring." *Current Biology* 24: 2855–60. https://doi.org/10.1016/j.cub.2014.10.039.

Fietz, Joanna, Hans Zischler, Claudia Schwiegk, Jürgen Tomiuk, Kathrin H. Dausmann, and Jörg U. Ganzhorn. 2000. "High Rates of Extra-Pair Young in the Pair-Living Fat-Tailed Dwarf Lemur, Cheirogaleus Medius." *Behavioral Ecology and Sociobiology* 49: 8–17. https://doi.org/10.1007/s002650000269.

Fox, Stephanie. 2015. "The Effect of Potential and Actual Paternity on Positive Male-Infant Behaviour in Ursine Colobus." Master's thesis, University of Calgary. https://doi.org/10.11575/PRISM/28090.

Fox, Stephanie et al. n.d. "Unpublished Paternity Data for Ursine Colobus."

Gatti, Silvia et al. n.d. "Unpublished Paternity Data for Western Lowland Gorillas Extracted from s. Gatti 2005 PhD Thesis."

Gerloff, Ulrike, Bianka Hartung, Barbara Fruth, Gottfried Hohmann, and Diethard Tautz. 1999. "Intracommunity Relationships, Dispersal Pattern and Paternity Success in a Wild Living Community of Bonobos (Pan Paniscus) Determined from DNA Analysis of Faecal Samples." *Proceedings of the Royal Society B* 266. https://doi.org/10.1098/rspb.1999.0762.

Gilby, Ian C., Lauren J. N. Brent, Emily E. Wroblewski, Rebecca S. Rudicell, Beatrice H. Hahn, Jane Goodall, and Anne E. Pusey. 2013. "Fitness Benefits of Coalitionary Aggression in Male Chimpanzees." *Behavioral Ecology and Sociobiology* 67: 373–81. https://doi.org/10.1007/s00265-012-1457-6.

Godoy, Irene, Linda Vigilant, and Susan E. Perry. 2016a. "Cues to Kinship and Close Relatedness During Infancy in White-Faced Capuchin Monkeys, Cebus Capucinus." *Animal Behaviour* 116: 139–51. https://doi.org/10.1016/j.anbehav.2016.03.031.

———. 2016b. "Inbreeding Risk, Avoidance and Costs in a Group-Living Primate, Cebus Capucinus." *Behavioral Ecology and Sociobiology* 70: 1601–11. https://doi.org/10.1007/s00265-016-2168-1.

Goossens, B., J. M. Setchell, S. S. James, S. M. Funk, L. Chikhi, A. Abulani, M. Ancrenaz, I. Lackman-Ancrenaz, and M. W. Bruford. 2006. "Philopatry and Reproductive Success in Bornean Orang-Utans (Pongo Pygmaeus)." *Molecular Ecology* 15: 2577–88. https://doi.org/10.1111/j.1365-294X.2006.02952.x.

Guo, Songtao, Weihong Ji, Ming Li, Hongli Chang, and Baoguo Li. 2010. "The Mating System of the Sichuan Snub-Nosed Monkey (Rhinopithecus Roxellana)." *American Journal of Primatology* 72: 25–32. https://doi.org/10.1002/ajp.20747.

Hayakawa, Sachiko. 2008. "Male–Female Mating Tactics and Paternity of Wild Japanese Macaques (Macaca Fuscata Yakui)." *American Journal of Primatology* 70: 986–89. https://doi.org/10.1002/ajp.20580.

Heistermann, Michael, Thomas Ziegler, Carel P. van Schaik, Kristin Launhardt, Paul Winkler, and J. Keith Hodges. 2001. "Loss of Oestrus, Concealed Ovulation and Paternity Confusion in Free-Ranging Hanuman Langurs." *Proceedings of the Royal Society B*. https://doi.org/10.1098/rspb.2001.1833.

Higham, James P., Michael Heistermann, Muhammad Agil, Dyah Perwitasari-Farajallah, Anja Widdig, and Antje Engelhardt. 2021. "Female Fertile Phase Synchrony, and Male Mating and Reproductive Skew, in the Crested Macaque." *Scientific Reports*. https://doi.org/10.1038/s41598-021-81163-1.

Huang, Xia, Nai-qing Hu, Colin A. Chapman, Kai He, Xue-long Jiang, Zhen-hua Guan, Peng-fei Fan, and Paul A. Garber. 2022. "Disassociation of Social and Sexual Partner Relationships in a Gibbon Population with Stable One-Male Two-Female Groups." *American Journal of Primatology*. https://doi.org/10.1002/ajp.23394.

Huchard, Elise, Alexandra Alvergne, Delphine Féjan, Leslie A. Knapp, Guy Cowlishaw, and Michel Raymond. 2010. "More Than Friends? Behavioural and Genetic Aspects of Heterosexual Associations in Wild Chacma Baboons." *Behavioral Ecology and Sociobiology* 64: 769–81. https://doi.org/10.1007/s00265-009-0894-3.

Huck, Maren, Eduardo Fernandez-Duque, Paul Babb, and Theodore Schurr. 2014. "Correlates of Genetic Monogamy in Socially Monogamous Mammals: Insights from Azara's Owl Monkeys." *Proceedings of the Royal Society B*. https://doi.org/10.1098/rspb.2014.0195.

Huck, Maren, Petra Löttker, Uta-Regina Böhle, and Eckhard W. Heymann. 2005. "Paternity and Kinship Patterns in Polyandrous Moustached Tamarins (Saguinus Mystax)." *American Journal of Physical Anthropology* 127: 449–64. https://doi.org/10.1002/ajpa.20136.

Inoue, Eiji, Etienne François Akomo-Okoue, Chieko Ando, Yuji Iwata, Mariko Judai, Shiho Fujita, Shun Hongo, Chimene Nze-Nkogue, Miho Inoue-Murayama, and Juichi Yamagiwa. 2013. "Male Genetic Structure and Paternity in Western Lowland Gorillas (Gorilla Gorilla Gorilla)." *American Journal of Physical Anthropology* 151: 583–88. https://doi.org/10.1002/ajpa.22312.

Inoue, Eiji, Miho Inoue-Murayama, Linda Vigilant, Osamu Takenaka, and Toshisada Nishida. 2008. "Relatedness in Wild Chimpanzees: Influence of Paternity, Male Philopatry, and Demographic Factors." *American Journal of Physical Anthropology* 137: 256–62. https://doi.org/10.1002/ajpa.20865.

Inoue, Eiji, and Osamu Takenaka. 2008. "The Effect of Male Tenure and Female Mate Choice on Paternity in Free-Ranging Japanese Macaques." *American Journal of Primatology* 70: 62–68. https://doi.org/10.1002/ajp.20457.

Ishizuka, Shintaro, Yoshi Kawamoto, Tetsuya Sakamaki, Nahoko Tokuyama, Kazuya Toda, Hiroki Okamura, and Takeshi Furuichi. 2018. "Paternity and Kin Structure Among Neighbouring Groups in Wild Bonobos at Wamba." *Royal Society Open Science* 5: 171006. https://doi.org/10.1098/rsos.171006.

Jack, Katharine M., and Linda M. Fedigan. 2006. "Why Be Alpha Male? Dominance and Reproductive Success in Wild White-Faced Capuchins (Cebus Capucinus)." In *New Perspectives in the Study of Mesoamerican Primates*, 367–86. Boston, MA: Springer. https://doi.org/10.1007/0-387-25872-8_18.

Jacobs, Rachel L., David C. Frankel, Riley J. Rice, Vera J. Kiefer, and Brenda J. Bradley. 2018. "Parentage Complexity in Socially Monogamous Lemurs (Eulemur Rubriventer): Integrating Genetic and Observational Data." *American Journal of Primatology*. https://doi.org/10.1002/ajp.22738.

Kappeler, Peter M., and Markus Port. 2008. "Mutual Tolerance or Reproductive Competition? Patterns of Reproductive Skew Among Male Redfronted Lemurs (Eulemur Fulvus Rufus)." *Behavioral Ecology and Sociobiology* 62: 1477–88. https://doi.org/10.1007/s00265-008-0577-5.

Kappeler, Peter M., and Livia Schäffler. 2008. "The Lemur Syndrome Unresolved: Extreme Male Reproductive Skew in Sifakas (Propithecus Verreauxi), a Sexually Monomorphic Primate with Female Dominance." *Behavioral Ecology and Sociobiology* 62: 1007–15. https://doi.org/10.1007/s00265-007-0528-6.

Keane, B., W. P. J. Dittus, and D. J. Melnick. 1997. "Paternity Assessment in Wild Groups of Toque Macaques Macaca Sinica at Polonnaruwa, Sri Lanka Using Molecular Markers." *Molecular Ecology* 6: 267–82. https://doi.org/10.1046/j.1365-294x.1997.00178.x.

Kenyon, Marina, Christian Roos, Vo Thanh Binh, and David Chivers. 2011. "Extrapair Paternity in Golden-Cheeked Gibbons (Nomascus Gabriellae) in the Secondary Lowland Forest of Cat Tien National Park, Vietnam." *Folia Primatologica* 82: 154–64. https://doi.org/10.1159/000333143.

Kümmerli, Rolf, and Robert D. Martin. 2005. "Male and Female Reproductive Success in Macaca Sylvanus in Gibraltar: No Evidence for Rank Dependence." *International Journal of Primatology* 26 (6): 1229–47. https://doi.org/10.1007/s10764-005-8851-0.

Langergraber, Kevin E., John C. Mitani, David P. Watts, and Linda Vigilant. 2013. "Male–Female Socio-Spatial Relationships and Reproduction in Wild Chimpanzees." *Behavioral Ecology and Sociobiology* 67: 861–73. https://doi.org/10.1007/s00265-013-1509-6.

Larney, Eileen. 2013. "The Influence of Genetic and Social Structure on Reproduction in Phayre's Leaf Monkeys (Trachypithecus Phayrei Crepusculus)." PhD thesis, Stony Brook University.

Launhardt, Kristin, Carola Borries, Cornelia Hardt, Jörg T. Epplen, and Paul Winkler. 2001. "Paternity Analysis of Alternative Male Reproductive Routes Among the Langurs (Semnopithecus Entellus) of Ramnagar." *Animal Behaviour* 61: 53–64. https://doi.org/10.1006/anbe.2000.1590.

Lawler, Richard R. 2007. "Fitness and Extra-Group Reproduction in Male Verreaux's Sifaka: An Analysis of Reproductive Success from 1989–1999." *American Journal of Physical Anthropology* 132: 267–77. https://doi.org/10.1002/ajpa.20507.

Lawler, Richard R., Alison F. Richard, and Margaret A. Riley. 2003. "Genetic Population Structure of the White Sifaka (Propithecus verreauxi verreauxi) at Beza Mahafaly Special Reserve, Southwest Madagascar (1992–2001)." *Molecular Ecology* 12: 2307–17. https://doi.org/10.1046/j.1365-294X.2003.01909.x.

Liu, Zhijin, Chengming Huang, Qihai Zhou, Youbang Li, Yuefeng Wang, Ming Li, Osamu Takenaka, and Akiko Takenaka. 2013. "Genetic Analysis of Group Composition and Relatedness in White-Headed Langurs." *Integrative Zoology* 8: 410–16. https://doi.org/10.1111/1749-4877.12048.

Masi, Shelly, Frédéric Austerlitz, Chloé Chabaud, Sophie Lafosse, Nina Marchi, Myriam Georges, Françoise Dessarps-Freichey, et al. 2021. "No Evidence for Female Kin Association, Indications for Extragroup Paternity, and Sex-Biased Dispersal Patterns in Wild Western Gorillas." *Ecology and Evolution*. https://doi.org/10.1002/ece3.7596.

McCarthy, M. S., J. D. Lester, M. Cibot, L. Vigilant, and M. R. McLennan. 2020. "Atypically High Reproductive Skew in a Small Wild Chimpanzee Community in a Human-Dominated Landscape." *Folia Primatologica* 91: 688–96. https://doi.org/10.1159/000508609.

Miller, Carrie M., Noah Snyder-Mackler, Nga Nguyen, Peter J. Fashing, Jenny Tung, Emily E. Wroblewski, Morgan L. Gustison, and Michael L. Wilson. 2021. "Extragroup Paternity in Gelada Monkeys, Theropithecus gelada, at Guassa, Ethiopia and a Comparison with Other Primates." *Animal Behaviour* 177: 277–301. https://doi.org/10.1016/j.anbehav.2021.05.008.

Minker, Mirjam M. I., Christopher Young, Federica Amici, Richard McFarland, Louise Barrett, J. Paul Grobler, S. Peter Henzi, and Anja Widdig. 2018. "Assessment of Male Reproductive Skew via Highly Polymorphic STR Markers in Wild Vervet Monkeys, Chlorocebus pygerythrus." *Journal of Heredity* 109: 780–90. https://doi.org/10.1093/jhered/esy048.

Moscovice, Liza R., Anthony Di Fiore, Catherine Crockford, Dawn M. Kitchen, Roman Wittig, Robert M. Seyfarth, and Dorothy L. Cheney. 2010. "Hedging Their Bets? Male and Female Chacma Baboons Form Friendships Based on Likelihood of Paternity." *Animal Behaviour* 79: 1007–15. https://doi.org/10.1016/j.anbehav.2010.01.013.

Moscovice, Liza, Robert Seyfarth, and Joan Silk. n.d. "Unpublished Paternity Data for Chacma Baboons."

Mouginot, Maud, Leveda Cheng, Michael L. Wilson, Joseph T. Feldblum, Veronika Städele, Emily E. Wroblewski, Linda Vigilant, et al. 2023. "Reproductive Inequality Among Males in the Genus Pan." *Philosophical Transactions of the Royal Society B* 378: 20220301. https://doi.org/10.1098/rstb.2022.0301.

Muniz, Laura, Susan Perry, Joseph H. Manson, Hannah Gilkenson, Julie Gros-Louis, and Linda Vigilant. 2010. "Male Dominance and Reproductive Success in Wild White-Faced Capuchins (Cebus capucinus) at Lomas Barbudal, Costa Rica." *American Journal of Primatology* 72: 1118–30. https://doi.org/10.1002/ajp.20876.

Newton-Fisher, Nicholas E., Melissa Emery Thompson, Vernon Reynolds, Christophe Boesch, and Linda Vigilant. 2010. "Paternity and Social Rank in Wild Chimpanzees (Pan troglodytes) from the Budongo Forest, Uganda." *American Journal of Physical Anthropology* 142: 417–28. https://doi.org/10.1002/ajpa.21241.

Nievergelt, Caroline M., Thomas Mutschler, Anna T. C. Feistner, and David S. Woodruff. 2002. "Social System of the Alaotran Gentle Lemur (Hapalemur griseus alaotrensis): Genetic Characterization of Group Composition and Mating System." *American Journal of Primatology* 57: 157–76. https://doi.org/10.1002/ajp.10046.

Nsubaga, Anthony M., Martha M. Robbins, Christophe Boesch, and Linda Vigilant. 2008. "Patterns of Paternity and Group Fission in Wild Multimale Mountain Gorilla Groups." *American Journal of Physical Anthropology* 135: 263–74. https://doi.org/10.1002/ajpa.20740.

Ohsawa, Hideyuki, Miho Inoue, and Osamu Takenaka. 1993. "Mating Strategy and Reproductive Success of Male Patas Monkeys (Erythrocebus patas)." *Primates* 34 (4): 533–44. https://doi.org/10.1007/BF02382664.

Oka, Teruki, and Osamu Takenaka. 2001. "Wild Gibbons' Parentage Tested by Non-Invasive DNA Sampling and PCR-Amplified Polymorphic Microsatellites." *Primates* 42 (1): 67–73. https://doi.org/10.1007/BF02640690.

Oklander, Luciana I., Martin Kowalewski, and Daniel Corach. 2014. "Male Reproductive Strategies in

Black and Gold Howler Monkeys (Alouatta Caraya).” *American Journal of Primatology* 76: 43–55. https://doi.org/10.1002/ajp.22191.

Parga, Joyce A., Michelle L. Sauther, Frank P. Cuozzo, Ibrahim Antho Youssouf Jacky, Richard R. Lawler, Robert W. Sussman, Lisa Gould, and Jennifer Pastorini. 2016. “Paternity in Wild Ring-Tailed Lemurs (Lemur Catta): Implications for Male Mating Strategies.” *American Journal of Primatology* 78 (12): 1316–25. https://doi.org/10.1002/ajp.22584.

Platner, Benjamin. 2005. “Das Genetische Paarungssystem von Lepilemur Ruficaudatus.” Master’s thesis, University of Zurich.

Pope, Theresa R. 1990. “The Reproductive Consequences of Male Cooperation in the Red Howler Monkey: Paternity Exclusion in Multi-Male and Single-Male Troops Using Genetic Markers.” *Behavioral Ecology and Sociobiology* 27 (6): 439–46. https://doi.org/10.1007/BF00164071.

Qi, Xiao-Guang, Cyril C. Grueter, Gu Fang, Peng-Zhen Huang, Jing Zhang, Yan-Mei Duan, Zhi-Pang Huang, Paul A. Garber, and Bao-Guo Li. 2020. “Multilevel Societies Facilitate Infanticide Avoidance Through Increased Extrapair Matings.” *Animal Behaviour* 161: 127–37. https://doi.org/10.1016/j.anbehav.2019.12.014.

Roberts, Su-Jen, Eleni Nikitopoulos, and Marina Cords. 2014. “Factors Affecting Low Resident Male Siring Success in One-Male Groups of Blue Monkeys.” *Behavioral Ecology* 25 (4): 852–61. https://doi.org/10.1093/beheco/aru060.

Rosenbaum, S., J. P. Hirwa, J. B. Silk, L. Vigilant, and T. S. Stoinski. 2015. “Male Rank, Not Paternity, Predicts Male–Immature Relationships in Mountain Gorillas, Gorilla Beringei Beringei.” *Animal Behaviour* 104: 13–24. https://doi.org/10.1016/j.anbehav.2015.02.025.

Ruiter, Jan R. de, Jan A. R. A. M. van Hooff, and Wolfgang Scheffrahn. 1994. “Social and Genetic Aspects of Paternity in Wild Long-Tailed Macaques (Macaca Fascicularis).” *Behaviour* 129 (3/4): 203–24. https://doi.org/10.1163/156853994X00613.

Schülke, Oliver, Peter M. Kappeler, and Hans Zischler. 2004. “Small Testes Size Despite High Extra-Pair Paternity in the Pair-Living Nocturnal Primate Phaner Furcifer.” *Behavioral Ecology and Sociobiology* 55: 293–301. https://doi.org/10.1007/s00265-003-0709-x.

Schwensow, Nina, Joanna Fietz, Kathrin Dausmann, and Simone Sommer. 2008. “MHC-Associated Mating Strategies and the Importance of Overall Genetic Diversity in an Obligate Pair-Living Primate.” *Evolutionary Ecology* 22: 617–36. https://doi.org/10.1007/s10682-007-9186-4.

Scott, Amy M., Graham L. Banes, Wuryantari Setiadi, Jessica R. Saragih, Tri Susanto Wahyu, Tatang Setia Mitra, and Cheryl D. Knott. 2024. “Flanged Males Have Higher Reproductive Success in a Completely Wild Orangutan Population.” *PLOS ONE* 19 (2): e0296688. https://doi.org/10.1371/journal.pone.0296688.

Silk, Joan et al. n.d. “Unpublished Paternity Data for Olive Baboons.”

Snyder-Mackler, Noah, Susan C. Alberts, and Thore J. Bergman. 2012. “Concessions of an Alpha Male? Cooperative Defence and Shared Reproduction in Multi-Male Primate Groups.” *Proceedings of the Royal Society B* 279: 3788–95. https://doi.org/10.1098/rspb.2012.0842.

Soltis, Joseph, Ruth Thomsen, and Osamu Takenaka. 2001. “The Interaction of Male and Female Reproductive Strategies and Paternity in Wild Japanese Macaques, Macaca Fuscata.” *Animal Behaviour* 62: 485–94. https://doi.org/10.1006/anbe.2001.1774.

Strier, Karen B., Paulo B. Chaves, Sérgio L. Mendes, Valéria Fagundes, and Anthony Di Fiore. 2011. “Low Paternity Skew and the Influence of Maternal Kin in an Egalitarian, Patrilocal Primate.” *Proceedings of the National Academy of Sciences* 108 (45): 17884–89. https://doi.org/10.1073/pnas.1116737108.

Sukmak, Manakorn, Worawidh Wajjwalku, Julia Ostner, and Oliver Schülke. 2014. “Dominance Rank, Female Reproductive Synchrony, and Male Reproductive Skew in Wild Assamese Macaques.” *Behavioral Ecology and Sociobiology* 68: 1097–1108. https://doi.org/10.1007/s00265-014-1721-z.

Surbeck, Martin, Kevin E. Langergraber, Barbara Fruth, Linda Vigilant, and Gottfried Hohmann. 2017. “Male Reproductive Skew Is Higher in Bonobos Than Chimpanzees.” *Current Biology* 27 (13): R503–5. https://doi.org/10.1016/j.cub.2017.05.039.

Tajima, Tomoyuki, Titol P. Malim, and Eiji Inoue. 2018. “Reproductive Success of Two Male Morphs in a Free-Ranging Population of Bornean Orangutans.” *Primates* 59: 127–33. https://doi.org/10.1007/s10329-017-0648-1.

Teichroeb, Julie A., Eva C. Wikberg, Nelson Ting, and Pascale Sicotte. 2014. “Factors Influencing Male

Affiliation and Coalitions in a Species with Male Dispersal and Intense Male–Male Competition, Colobus Vellerosus." *Behaviour* 151 (7): 1045–66. https://doi.org/10.1163/1568539X-00003089.

Trébouet, Florian. 2019. "Male Reproductive Strategies in Wild Northern Pig-Tailed Macaques (Macaca Leonina): Testing the Priority-of-Access Model." PhD thesis, Southern Illinois University Carbondale.

Utami, Sri Suci, Benoît Goossens, Michael W. Bruford, Jan R. de Ruiter, and Jan A. R. A. M. van Hooff. 2002. "Male Bimaturism and Reproductive Success in Sumatran Orang-Utans." *Behavioral Ecology* 13 (5): 643–52. https://doi.org/10.1093/beheco/13.5.643.

Van Belle, Sarie, E. Martins, and A. Di Fiore. 2016. "Patterns of Paternity in Wild Socially-Monogamous Titis (Callicebus Discolor) and Sakis (Pithecia Aequatorialis) at the Tiputini Biodiversity Station, Ecuador (Conference Abstract)." In *International Journal of Primatology and American Society of Primatologists.*

Vigilant, Linda, Michael Hofreiter, Heike Siedel, and Christophe Boesch. 2001. "Paternity and Relatedness in Wild Chimpanzee Communities." *Proceedings of the National Academy of Sciences* 98 (23): 12890–95. https://doi.org/10.1073/pnas.231320498.

Vigilant, Linda, Justin Roy, Brenda J. Bradley, Colin J. Stoneking, Martha M. Robbins, and Tara S. Stoinski. 2015. "Reproductive Competition and Inbreeding Avoidance in a Primate Species with Habitual Female Dispersal." *Behavioral Ecology and Sociobiology* 69: 1163–72. https://doi.org/10.1007/s00265-015-1930-0.

Walker, Kara K., Rebecca S. Rudicell, Yingying Li, Beatrice H. Hahn, Emily Wroblewski, and Anne E. Pusey. 2017. "Chimpanzees Breed with Genetically Dissimilar Mates." *Royal Society Open Science* 4: 160422. https://doi.org/10.1098/rsos.160422.

Wang, Bai-Shi, Zhen-Long Wang, Jun-Dong Tian, Zhen-Wei Cui, and Ji-Qi Lu. 2015. "Establishment of a Microsatellite Set for Noninvasive Paternity Testing in Free-Ranging Macaca Mulatta Tcheliensis in Mount Taihangshan Area, Jiyuan, China." *Zoological Studies* 54: 8. https://doi.org/10.1186/s40555-014-0100-9.

Weiss, Alexander, Joseph T. Feldblum, Drew M. Altschul, David Anthony Collins, Shadrack Kamenya, Deus Mjungu, Steffen Foerster, Ian C. Gilby, Michael L. Wilson, and Anne E. Pusey. 2023. "Personality Traits, Rank Attainment, and Siring Success Throughout the Lives of Male Chimpanzees of Gombe National Park." *PeerJ* 11: e15083. https://doi.org/10.7717/peerj.15083.

Wikberg, Eva C., Katharine M. Jack, Linda M. Fedigan, Fernando A. Campos, Akiko S. Yashima, Mackenzie L. Bergstrom, Tomohide Hiwatashi, and Shoji Kawamura. 2017. "Inbreeding Avoidance and Female Mate Choice Shape Reproductive Skew in Capuchin Monkeys (Cebus Capucinus Imitator)." *Molecular Ecology* 26: 653–67. https://doi.org/10.1111/mec.13898.

Wimmer, Barbara, and Peter M. Kappeler. 2002. "The Effects of Sexual Selection and Life History on the Genetic Structure of Redfronted Lemur, Eulemur Fulvus Rufus, Groups." *Animal Behaviour* 64: 557–68. https://doi.org/10.1006/anbe.2002.4003.

Wroblewski, Emily E., Carson M. Murray, Brandon F. Keele, Joann C. Schumacher-Stankey, Beatrice H. Hahn, and Anne E. Pusey. 2009. "Male Dominance Rank and Reproductive Success in Chimpanzees, Pan Troglodytes Schweinfurthii." *Animal Behaviour* 77 (4): 873–85. https://doi.org/10.1016/j.anbehav.2008.12.014.

Wu, Fan, Jia Liu, Derek W. Dunn, Yixin Shang, Shiyu Jin, Huihui Du, Yuanchun Wu, et al. 2025. "Beyond the Alpha: Extra-Pair Paternities and Male Reproductive Success in a Primate Multilevel Society." *Ecology and Evolution* 15: e70928. https://doi.org/10.1002/ece3.71749.

Xiang, Zuo-Fu, Bang-He Yang, Yang Yu, Hui Yao, Cyril C. Grueter, Paul A. Garber, and Ming Li. 2014. "Males Collectively Defend Their One-Male Units Against Bachelor Males in a Multi-Level Primate Society." *American Journal of Primatology* 76: 609–17. https://doi.org/10.1002/ajp.22254.

Yamane, A., T. Shotake, A. Mori, A. I. Boug, and T. Iwamoto. 2003. "Extra-Unit Paternity of Hamadryas Baboons (Papio Hamadryas) in Saudi Arabia." *Ethology Ecology and Evolution* 15 (4): 379–87. https://doi.org/10.1080/08927014.2003.9522664.

Yang, Bang-He, Baoping Ren, Zuo-Fu Xiang, Jingyuan Yang, Hui Yao, Paul A. Garber, and Ming Li. 2014. "MHC-DRB Exon II Variation in a Wild Population of Rhinopithecus Roxellana." *Integrative Zoology* 9: 598–612. https://doi.org/10.1111/1749-4877.12084.
